# Deep brain stimulation in the subthalamic nucleus for Parkinson’s disease can restore dynamics of striatal networks

**DOI:** 10.1101/2021.08.29.458121

**Authors:** Elie M Adam, Emery N Brown, Nancy Kopell, Michelle M McCarthy

**Affiliations:** Picower Institute for Learning and Memory, Massachusetts Institute of Technology, Cambridge, MA; Department of Anesthesia, Critical Care and Pain Medicine, Massachusetts General Hospital, Boston, MA; Department of Mathematics and Statistics, Boston University, Boston, MA

**Author notes:** N.K. and M.M. contributed equally to this work.

**Keywords:** Deep brain stimulation, Rhythms and oscillations, Parkinson’s disease, Striatum, Subthalamic nucleus

## Abstract

Deep brain stimulation (DBS) of the subthalamic nucleus (STN) is highly effective in alleviating movement disability in patients with Parkinson’s disease (PD). However, its therapeutic mechanism of action is unknown. The healthy striatum exhibits rich dynamics resulting from an interaction of beta, gamma and theta oscillations. These rhythms are at the heart of selection, initiation and execution of motor programs, and their loss or exaggeration due to dopamine (DA) depletion in PD is a major source of the behavioral deficits observed in PD patients. Interrupting abnormal rhythms and restoring the interaction of rhythms as observed in the healthy striatum may then be instrumental in the therapeutic action of DBS. We develop a biophysical networked model of a BG pathway to study how abnormal beta oscillations can emerge throughout the BG in PD, and how DBS can restore normal beta, gamma and theta striatal rhythms. Our model incorporates STN projections to the striatum, long known but understudied, that were recently shown to preferentially target fast spiking interneurons (FSI) in the striatum. We find that DBS in STN is able to normalize striatal medium spiny neuron (MSN) activity by recruiting FSI dynamics, and restoring the inhibitory potency of FSIs observed in normal condition. We also find that DBS allows the re-expression of gamma and theta rhythms, thought to be dependent on high DA levels and thus lost in PD, through cortical noise control. Our study shows how BG connectivity can amplify beta oscillations, and delineates the role of DBS in disrupting beta oscillations and providing corrective input to STN efferents to restore healthy striatal dynamics. It also suggests how gamma oscillations can be leveraged to enhance or supplement DBS treatment and improve its effectiveness.

## Introduction

Deep brain stimulation (DBS) in the subthalamic nucleus (STN), the sole excitatory nucleus of the basal ganglia, elicits a remarkable effect of rapidly restoring to almost normal, the very disabling motor symptoms of Parkinson’s Disease (PD). However, the mechanism of DBS efficacy remains a mystery. It is generally thought that DBS works though its systems-level effects on networks within and between the nuclei of the basal ganglia (BG), thalamus and cortex [1]. The motor symptoms of bradykinesia and rigidity are correlated with exaggerated beta frequency (~ 15-30Hz) oscillations in STN local field potential (LFP) in PD patients [2, 3]. Suppression of beta oscillations following high frequency DBS to STN correlates with augmentation of motor function in PD patients [4]. This suggests that some of the efficacy of high frequency DBS in STN for PD symptoms may lie in the ability of DBS to reduce the pathologically elevated beta oscillations within the corticobasal ganglia-thalamic (CBT) loop. Indeed, models have proposed a mechanistic role for DBS in STN in disrupting the propagation of aberrant oscillations to STN efferents [5] and normalizing output nuclei of the BG [6, 7]. This normalization is found essential to restore relay reliability in the thalamus, which modeling suggests goes awry in Parkinsonian conditions due to abnormal BG output [8–11]. Restoring thalamic reliability is likely an outcome of network interactions following DBS in STN, and it has been suggested that DBS engages a mechanism of converging network-wide input onto the striatum to achieve regularity of firing at the output of the BG [12]. However, previous modeling work largely put the emphasis of the effects of DBS on the BG output, and ignored its effect on internal BG dynamics. While restoring normal brain function indeed necessitates restoring reliable thalamocortical relay, matching the intricacies of action selection and voluntary motor control further requires the richness of the dynamics normally observed inside the BG nuclei.

Specifically, previous work from our group has shown that increased excitability in striatal medium spiny neurons (MSNs) expressing D2-recepters, which results from loss of dopamine (DA), increases beta oscillations in striatal networks [13] (See Discussion for the role of striatum in creating pathological beta). These beta oscillations are generated from inhibitory MSN interactions, in the presence of high cholinergic tone during PD. Thus, a DBS mechanism effective at restoring BG function, and more particularly striatal function, needs to be capable of rectifying this source of aberrant beta activity. No such mechanism has yet been studied nor proposed. Furthermore, beta [14–17], gamma (≳40Hz) [18, 19] and theta (~4-8Hz) oscillations [20, 21] are normally expressed in a DA-dependent manner in striatal networks to drive behavior, and the loss of DA in PD can disrupt the formation of these rhythms, as later shown. However, their coexistence may be necessary for normal behavioral function. The question we seek to answer is how striatal network level dynamics are restored, through DBS in STN, despite persisting cellular level disruptions due to loss of DA. To answer this question, building on established computational models of striatum [13, 22], we explore network activity associated with a previously understudied but direct connection from STN to striatum [23–28]. Recent research shows that the direct STN to striatum pathway projects strongly and almost exclusively to striatal fast spiking interneurons (FSIs) [28].

There is uncertainty about the effect of DBS on STN and its efferents [29, 30]. The major experimental evidence is that high frequency stimulation (HFS) (≳100Hz) of STN suppresses somatic activity through depolarization blockade [31, 32] and stimulates axonal terminals at a high frequency [33, 34]. We modeled DBS in STN as a combination of these two aspects: DBS suppresses STN somatic activity, thereby suppressing the beta activity in STN, and replaces it with HFS at the level of STN axons. Our simulations show that HFS of the STN-FSI pathway fully restores striatal network dynamics including a reduction of beta and, most surprisingly, the re-expression of gamma and theta, striatal rhythms previously thought to be dependent on high levels of DA [22, 35–37]. Moreover, our models suggest that the gamma/theta and beta dynamics can be modulated by cortical input during DBS, rather than by DA, thus allowing an alternative mechanism to DA modulation during task performance. The diverse functions of striatal networks are still elusive, but the network dynamics that underlie the possible functions have been widely reported and characterized [14–21]. We thus confine our study of function restoration to an analysis of restoration of striatal dynamics, and leave the complexity of how these dynamics enable various functions to further investigation. Our study further highlights how the parkinsonian STN can amplify beta in striatum and thus, throughout the CBT loop, via this direct feedback pathway to striatum. We find that DBS not only normalizes striatum but also interferes with this amplifying feedback loop by its dampening effect on STN somatic activity.

## Results

### BG dynamics in normal conditions show intrinsically generated beta, gamma and theta oscillations

To study striatal dynamics, we developed a biophysical networked model of interacting neuronal populations in the BG, and studied it in different conditions: normal, parkinsonian and parkinsonian with DBS. DA levels fluctuate in normal conditions with effects on striatal dynamics. We will refer to baseline condition as the normal condition with baseline levels of DA, and separately introduce a normal condition with high levels of DA.

The core of our model consisted of a striatal population of medium spiny neurons (MSNs) inhibited by fast spiking interneurons (FSIs). We further modeled a population of subthalamic nucleus (STN) neurons projecting sparsely to the FSIs (Fig 1A), and completed a loop in the BG by modeling a population of external globus pallidus (GPe) neurons that project to STN and receive MSN projections (Fig 1A). This MSN-GPe-STN-FSI-MSN loop defines a neural pathway that partially goes through the indirect pathway of BG and offers the anatomical substrate for beta oscillations to be sustained and amplified, in PD. In baseline conditions, MSNs fire at an average rate (mean±SD) of 1.21±0.08 spk.s^−1^ (Fig 1B). Overall, our model shows an absence of beta oscillations throughout the network, as evidenced by the spectra of the four populations (Fig 1C). However, an isolated MSN network in baseline conditions does exhibit weak low-beta activity [13] (See Supplementary Information **SI**.A.1 and Figure S1A–C for details). But, in the core striatal model, we find that FSIs exhibit sparse gamma oscillations [22] (See **SI**.A.2 and Figure S1D−F for details) that inhibit MSN beta activity (See **SI**.A.3 and Figure S2A for details). The behavior of the core striatal model remains unaltered when connected to the greater network comprising GPe and STN (See **SI**.A.4 for details). The generation of beta in the MSNs is immediately suppressed by the FSIs and is not allowed to propagate throughout the loop.

**Figure 1:**
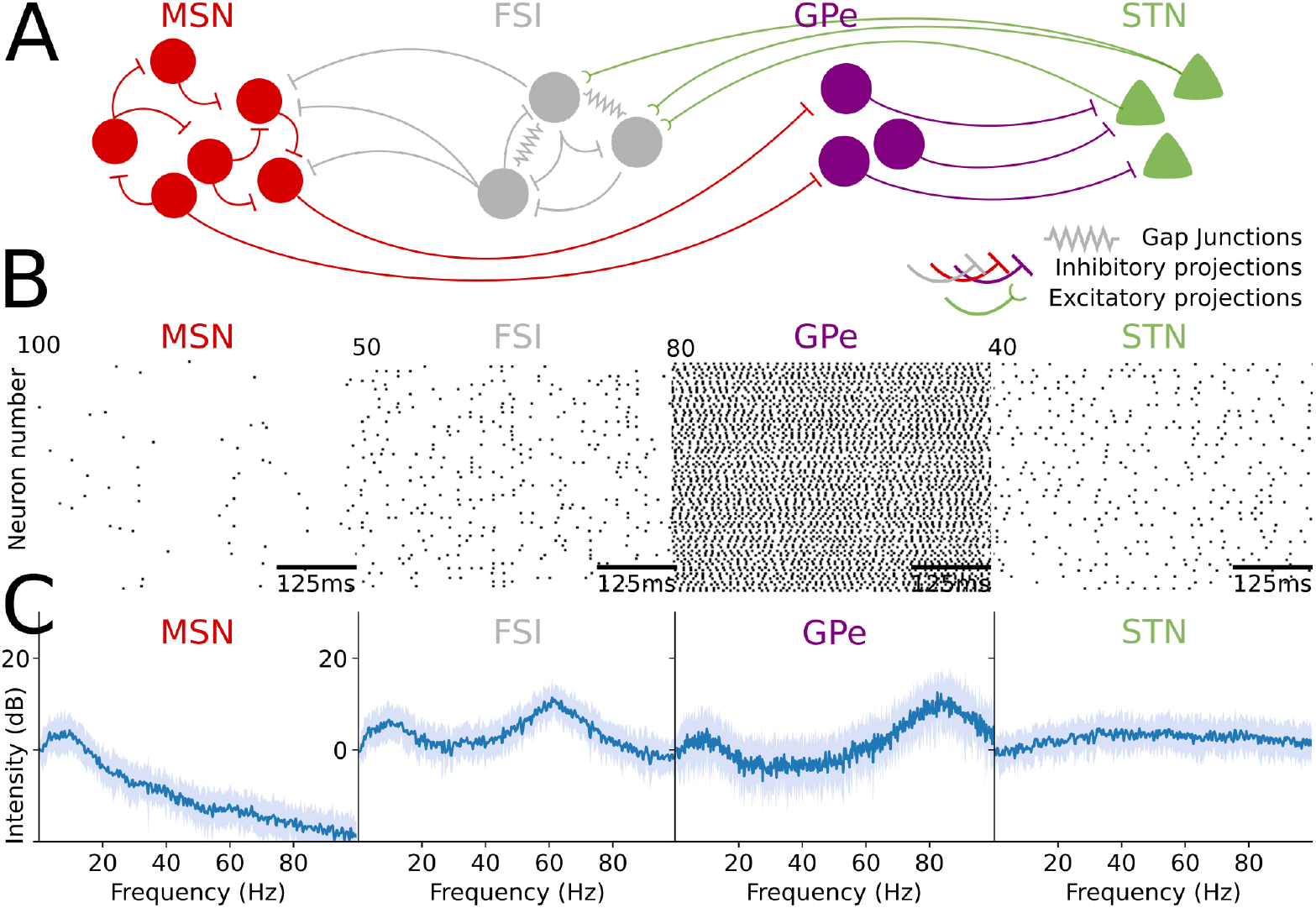
Population dynamics in baseline condition. **(A)** Schematic illustrating the network structure of the biophysical neuronal model, comprised of MSN, FSI, GPe and STN neurons. **(B)** Raster plots showing spiking activity of MSN, FSI, GPe and STN neurons, all in baseline condition (MSN firing rate: mean±SD=1.21 ± 0.08 spk.s^−1^, N=25 simulations). **(C)** Graphs showing the average (blue) and standard deviation (light blue) of the spectra of the MSN, FSI, GPe and STN population activity, all in baseline condition (N=25 simulations).

Nevertheless, beta oscillations in normal condition do appear in MSNs, and the BG more generally, through bursts of beta activity [38–40]. At high DA levels, the FSI excitability and gap junction conductance are increased, pushing the FSIs to spike at nested theta/gamma oscillations (Fig S3A). FSI activity then comprises synchronized bursts of gamma activity, interleaved with quiescence, appearing at theta cycles, observed in spiking activity and in spectral content showing peaks at theta (7.27±0.46Hz) and gamma frequencies (77.23±3.14Hz). The bursts of gamma activity have the ability to momentarily suppress the MSN network, leaving it to rebound during the FSI quiescence period and generate burst of beta oscillations [22]. This activity mode emerges from the interactions in our core striatal model (Fig S3B−D) and remains unaltered when connected to the greater network comprising GPe and STN. Synchronization of FSI activity into nested theta/gamma cycles is at the heart of enabling switches in cell-assemblies representing different motor programs [18, 41], with beta oscillations sustaining cell assemblies [20, 42]. We detail these results in later sections showing how DBS recovers such dynamics that are lost in PD.

### Loss of dopamine in PD leads to resonating beta activity throughout the BG

In PD, there are two effects that we included in our model: the loss of DA [43, 44] and the upregulation of striatal cholinergic levels as a result of DA depletion [45–47]. We modeled the changes in dopaminergic and cholinergic level through biophysical perturbations (Fig 2A). We find that these changes result in an increase in beta frequency power in the striatum, the GPe and the STN, consistent with what is observed clinically [13, 48–50].

**Figure 2:**
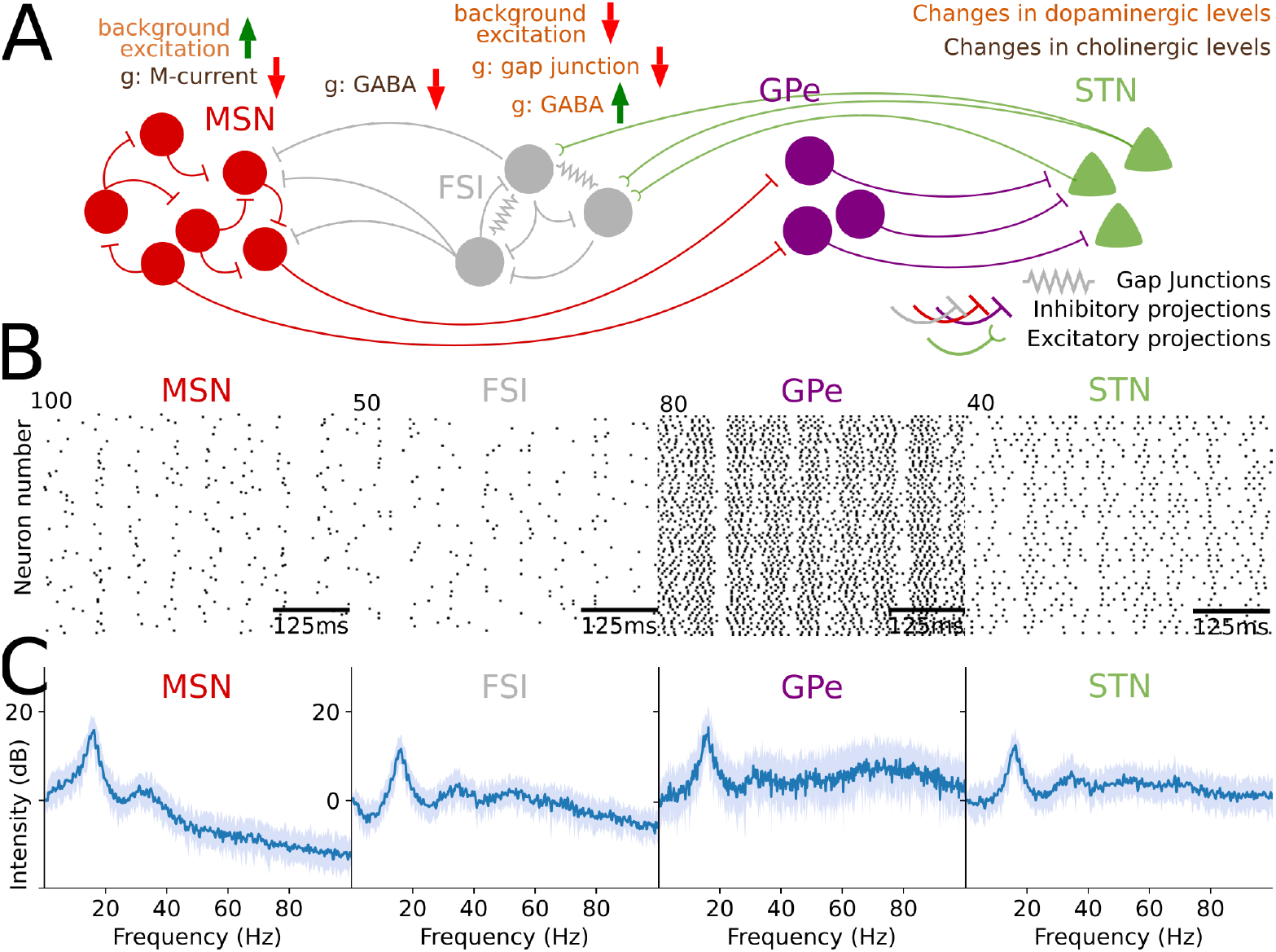
Population dynamics in PD. **(A)** Schematic illustrating the parametric changes in the biophysical model in PD from parameters in baseline condition. The loss of DA is modeled in multiple ways: (i) an increase of background excitation onto the MSNs expressing D2 receptors [45], (ii) a decrease in background excitation for FSIs [51] and (iii) changes in effective connectivity among the FSIs (decreased electrical conductance for the gap junctions [52], and increase in their interneuronal GABAa maximal conductance [51]). The increase in cholinergic tone (i) decreases the maximal MSN M-current conductance (via ACh action on M1 receptors [45]), and (ii) decreases the inhibitory maximal conductance from FSIs to MSNs [53]. **(B)** Raster plots showing spiking activity of MSN, FSI, GPe and STN neurons, all in PD (MSN firing rate: mean±SD=4.85±0.13 spk.s^−1^, N=25 simulations). **(C)** Graphs showing the average (blue) and standard deviation (light blue) of the spectra of the MSN, FSI, GPe and STN population activity, all in PD (N=25 simulations).

Our simulations show that an isolated D2 MSN network (Fig S4A−C, S5A–C) and our core striatal model (Fig S5D–F) produce beta oscillations during PD (See **SI**.A.5 for details). Embedding our core striatal model into the closed loop (Fig 2A), to include STN and GPe, amplifies beta-band activity during PD. We observe beta-band activity in all four populations MSN, FSI, STN and GPe, directly through spiking activity (Fig 2B) and through spectral power peaks in the beta-band (at 15.80 ± 0.43 Hz in MSNs) (Fig 2C) appearing in all four populations. The closed-loop system further creates resonance at beta frequencies as evidenced by additional power at harmonics, appearing in the spectra of population activity (Fig 2C) (See **SI**.A.6 for details). It is the increase of excitability of the MSNs that generates beta activity in the striatum (Fig S6A–D) (See **SI**.A.7 for details). This activity reaches the weakened FSIs, through the STN and GPe (Fig 2B), which in turns patterns the FSIs at beta frequencies (Fig 2C). Instead of FSIs suppressing MSN beta activity via their sparse gamma, FSIs become a conduit for beta-activity: the MSNs will resonate to the FSI beta-inhibition due to their intrinsic and network dynamics, and beta activity is then amplified in MSNs and throughout the loop (See **SI**.A.8 for details, and Figures S7A–C and S8A,B for illustrations on resonance properties).

Our modeling suggests an instrinsic striatal origin of beta activity that is propagated throughout the BG loop in PD conditions, but we find a similar amplification of exogeneous beta under PD conditions. We investigated the effects of adding an exogenous beta activity directly into STN to model the two alternative sources. We find that an additional exogenous input into STN amplifies the existing beta oscillations throughout the loop if provided at the resonating frequency (Fig S9A,B) and entrains the BG oscillations if not (Fig S9C–F) (See **SI**.A.9 for details). These results are consistent with experimental findings showing BG activity phase-locking to cortical beta bursts in normal and parkinsonian conditions [40]. Adding an exogenous beta input to MSNs instead of STN during PD also produces similar phenomena (Fig S10A–F).

### DBS in STN during PD can normalize MSN activity by restoring effective FSI inhibition

We modeled DBS in STN as a combination of two effects, considered in [30]: DBS suppresses somatic activity, thereby suppressing the beta activity derived from STN, and replaces it with high frequency activity at the level of the axons (Fig 3A). Essentially, DBS has the capability of decoupling STN axonal activity from STN somatic activity, thereby breaking beta oscillations in the BG loop and offering network excitation through HFS.

**Figure 3:**
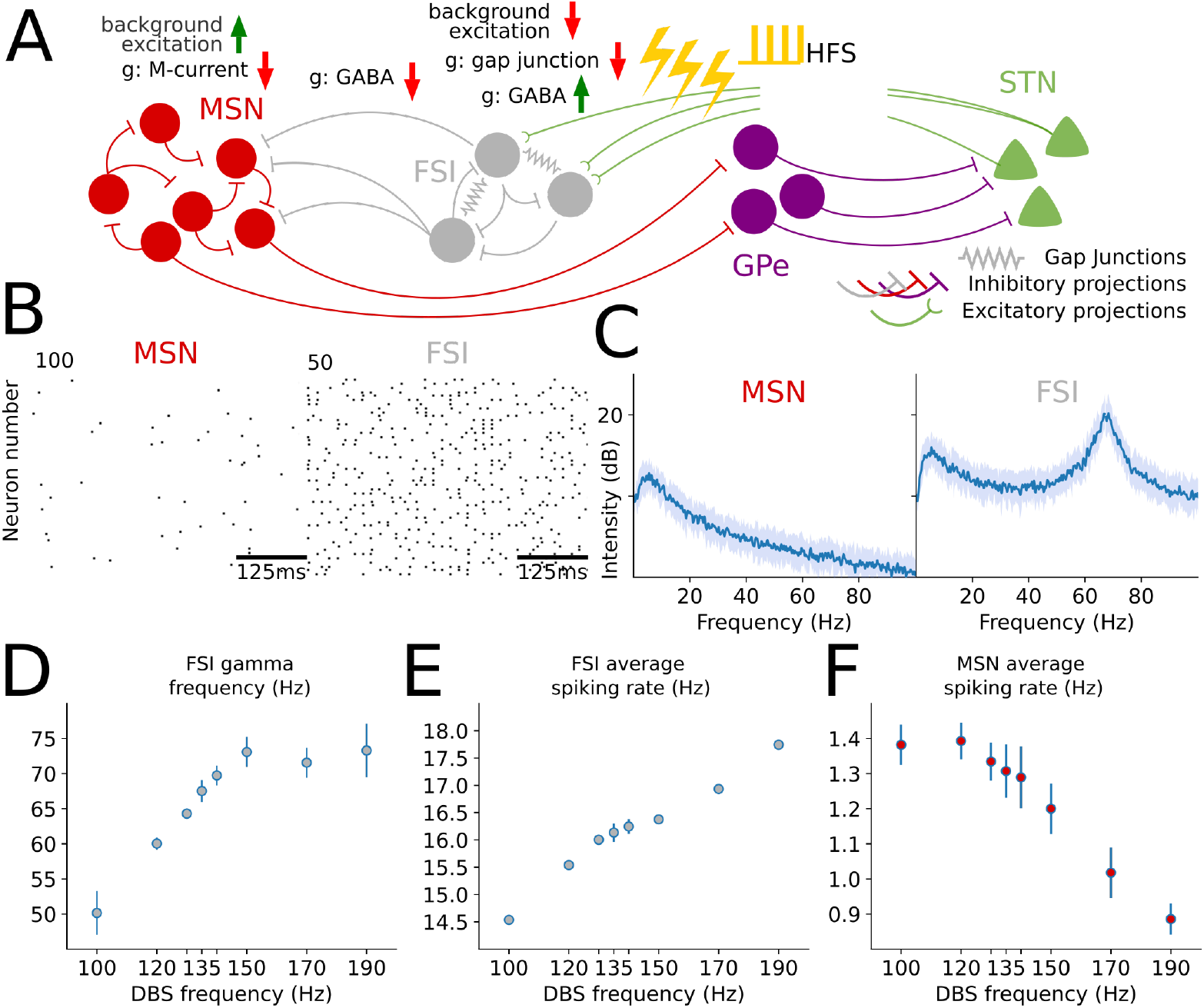
Population dynamics during DBS in PD. **(A)** Schematic illustrating the parametric changes in the biophysical model in PD during DBS from parameters in baseline condition. **(B)** Raster plots showing spiking activity of MSN and FSI neurons, all in PD during DBS (MSN firing rate: mean±SD=1.31 ± 0.07 spk.s^−1^, N=25 simulations). **(C)** Graphs showing the average (blue) and standard deviation (light blue) of the spectra of the MSN and FSI population activity, all in PD during DBS. **(D)** Graph showing the average FSI gamma oscillation frequency (bars representing standard deviation) as a function of DBS frequency (10 simulations per simulated frequency). **(E)** Graph similar to (D) showing the average FSI firing rate as a function of DBS frequency. **(F)** Graph similar to (D) showing the average MSN firing rate as a function of DBS frequency.

We modeled HFS by a train of voltage pulses subject to three parameters: (i) stimulation frequency, (ii) stimulation voltage and (iii) pulse width. We fixed the applied voltage, and varied the remaining two parameters: stimulation frequency and pulse width (See **SI**.A.10 for details).

We find that DBS at 135Hz with 150*μs* pulse-width during parkinsonian conditions normalizes MSN activity, lowering its firing rate to 1.31 ± 0.07 spk.s^−1^ similar to what is observed in baseline conditions (Fig 3B), and restoring its spectral properties to what is found in baseline conditions by removing the aberrant beta oscillations (Fig 3C). We find that DBS achieves MSN activity normalization by altering FSI activity: the level of excitation in FSIs is increased, leading to higher spiking rates that include bursting at gamma frequencies (Fig 3B, C) and that the gamma frequency of the FSI population follows around half of the stimulation frequency (at 67.1 ± 0.85Hz for stimulation at 135Hz) (corresponding to the peak in Fig 3C).

We next find that FSIs can play a decisive role in determining the optimal frequency of DBS. As we vary the stimulation frequency, we find that the FSIs produce bursts at half of the stimulation frequency (Fig 3D) (See **SI**.A.11 and Figure S11A for details). Overall, increasing the DBS frequency also increases FSI firing rate (Fig 3E) and thereby increasing FSI-MSN inhibition, leading to a decrease in MSN firing rate (Fig 3F) (See **SI**.A.12 and Figure S11B,C for details). Thus, our model predicts that the optimal frequency for DBS will depend on the average MSN spiking rate in PD, a value which will vary by patient. Increasing the pulse-width has a similar effect (See **SI**.A.13 and Figure S11D−F for details).

In lower frequency ranges, 60Hz and upwards, the FSIs oscillate at the DBS frequency (Fig S12A,B: an example for DBS at 65Hz), thereby justifying clinical improvement at these low frequencies too [54, 55] (Fig S12A, average MSN firing rate at 65Hz: 1.65 ± 0.08 spk.s^−1^). However, we will show in a later section that such low frequencies fail sustain theta/gamma FSI oscillations in PD conditions observed in normal dynamics under high levels of DA. Furthermore, stimulation at beta frequencies result in increased beta oscillation in MSNs (Fig S12C,D) (Average MSN firing rate at 5.57 ± 0.23 spk.s^−1^, greater than that of MSNs in PD condition without DBS, p¡0.001, Welch’s t-test), which is consistent with findings showing worsening of symptoms [54, 56, 57].

### DBS recovers theta and gamma oscillations lost during PD

At high DA levels in normal condition, FSI activity comprises synchronized bursts of gamma activity, interleaved with quiescence, appearing at theta cycles [22], clearly observed from the spiking activity (Fig 4A) and the spectral content showing peaks at theta (7.27 ± 0.46Hz) and gamma frequencies (77.23 ± 3.14Hz) (Fig 4B). The loss of DA during PD renders the FSIs unable to achieve such a dynamical state. This is partly due to a decreased excitation and an inability to synchronize in the presence of high beta noise with weakened gap junctions (See **SI**.A.14 for details). However, we find that DBS increases FSI excitability, and disrupts the beta noise coming into FSIs. This provides the opportunity for gamma bursts to arise, and that opportunity is governed by how well they are able to synchronize due to background noise. Indeed, background noise that is highly correlated among FSIs allows the FSIs to achieve synchrony, thereby replicating what is observed in conditions of high levels of DA, at the level of spiking activity (Fig 4C) and spectral content (Fig 4D) (gamma frequency at 66.81 ± 3.68 Hz and theta frequency at 6.11 ± 0.68 Hz). Background noise that is highly uncorrelated among FSIs would break synchrony, and thus would replicate the conditions we observed in baseline normal condition (Fig 1B,C). This suggests that DBS is able to substitute the mechanism of high/baseline DA, lost in PD, by a mechanism of correlated/uncorrelated noise to restore FSI and MSN dynamics.

**Figure 4:**
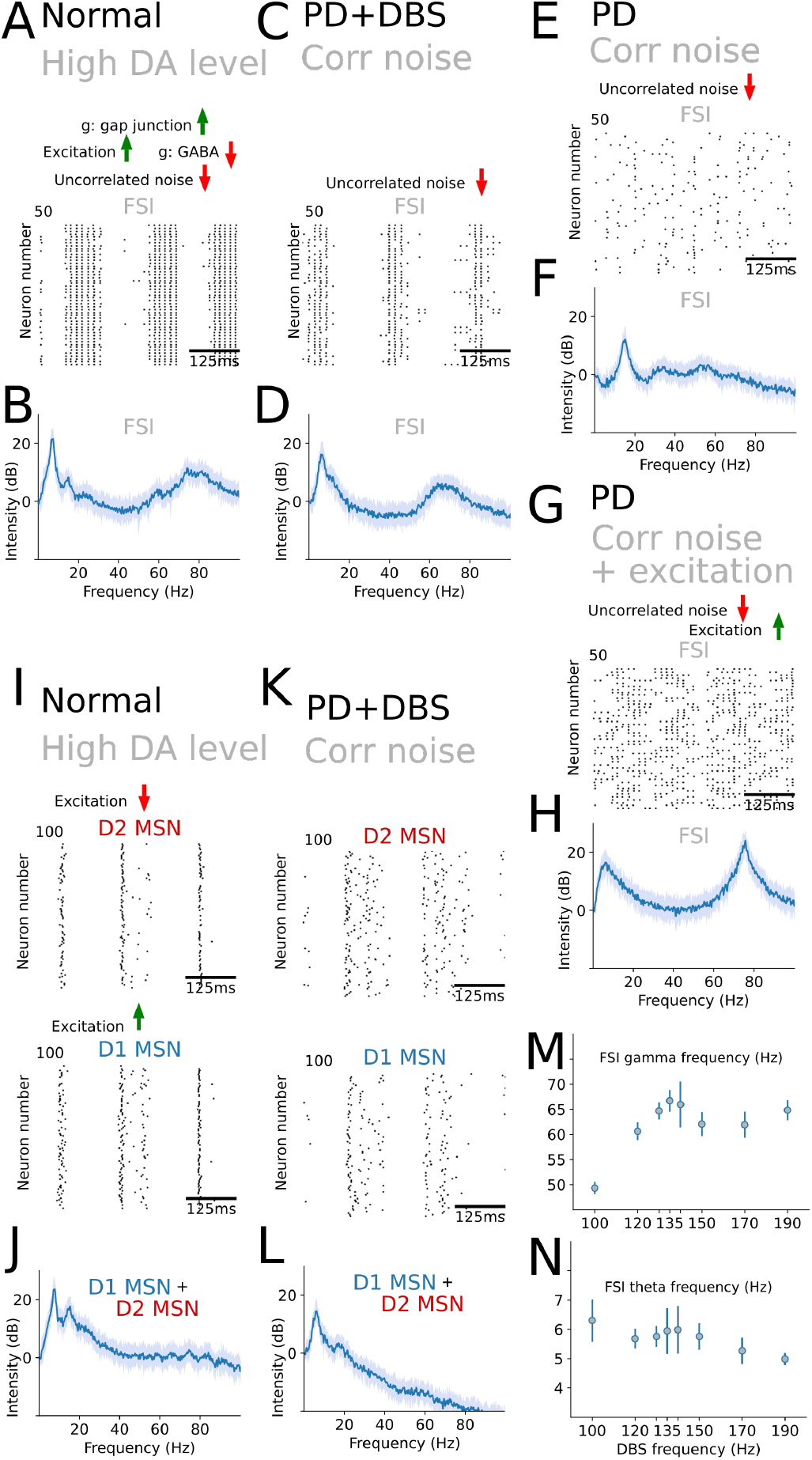
DBS can restore the DA functionality lost during PD. **(A)** Raster plots showing spiking activity of FSI neurons, in normal conditions with high level of DA. **(B)** Graph showing the average (blue) and standard deviation (light blue) of the spectrum of FSI population activity, in normal conditions with high level of DA. **(C)**, **(D)** Graphs displaying population activity of FSI neurons, as in (A) and (B), but for PD with DBS and a synchronized noise regime. **(E)** Raster plots showing spiking activity of FSI neurons, in PD within a synchronized noise regime. **(F)** Graph showing the average (blue) and standard deviation (light blue) of the spectrum of FSI population activity, in PD within a correlated noise regime. **(G)**, **(H)** Graphs displaying population activity of FSI neurons, as in (E) and (F), but for PD within a correlated noise regime and added excitation for the FSIs to drive them individually at a theta/gamma oscillation. **(I)**, **(J)** Graphs displaying population activity of MSN neurons, as in (A) and (B), for normal conditions with high level of dopamine. The raster plots show D1 and D2 MSN activity separately. **(K)**, **(L)** Graphs displaying population activity of MSN neurons, as in (I) and (J), but for PD with DBS and a correlated noise regime. **(M)** Graph showing the average FSI gamma oscillation frequency (bars representing standard deviation) as a function of DBS frequency (10 simulations per simulated frequency). **(N)** Graph similar to (M) showing the average FSI theta oscillation frequency as function of DBS frequency. All average spectra are derived from 25 simulations in each condition.

We modeled the source of uncorrelated/correlated background noise to be coming from cortical input. We do not expect that mechanism of switching correlation to be one generated only in PD. Per our interpretation, in a regime of changing motor plans, we expect cortex to be engaged, with its activity correlated and conveyed to the striatum. This setting can coincide with high DA levels, and deliver noise correlated among FSIs in normal conditions. In other regimes coinciding with baseline DA levels, cortical activity is uncorrelated and only conveys uncorrelated noise onto the FSIs. In normal conditions, however, the gap junctions are strong enough to overcome the noise whether it is uncorrelated or correlated. Therefore, although that functionality is always present, it need not be effectively used by the FSIs and can gain an effective use in PD during DBS, where DA levels are pathologically low. It also may be partially redundant in normal condition: correlated noise promotes more synchrony among FSIs in normal condition, especially if the level of DA is low for some normal reason. However, the correlated noise by itself cannot bring the FSIs to robustly spike a theta/gamma without additional excitation onto the FSIs. This excitation is provided by a high level of DA in normal conditions and by DBS in PD, but can also be provided by phasic activity from thalamic/cortical projections.

Importantly, in PD conditions without DBS, it is not possible to recover synchronized theta/gamma oscillations in FSI population activity with correlated noise (Fig 4E−F), even if we input additional excitation (Fig 4G−H) (See **SI**.A.15 for details).

### DBS recovers beta bursts lost during PD

At the level of MSNs during high DA conditions, the activity of D1 MSNs is increased, and D1 MSNs become a key player in the direct pathway of the BG. The activity of D2 MSNs is oppositely decreased. To study the effect of DBS on recreating MSN dynamics appearing in normal conditions with high DA, we then additionally modeled a population of D1 MSNs in our network. We modeled them to receive FSI projections, but they do not project to the GPe, as they are not considered canonically to be part of the indirect pathway of the BG. We also assumed that D1 and D2 MSNs are not interconnected by GABAa channels, to keep the role of D2 MSNs and their effect in our modeled indirect pathway loop unaltered.

In our simulations for normal conditions with high levels of DA, we found that both D1 and D2 MSNs were spiking when, and only when, FSIs were in their off cycle (Fig 4I). These bursts are produced at 7.21 ± 0.51Hz theta cycles (Fig 4I), and exhibit beta oscillations (peak at 15.72 ± 0.72Hz) in the short bursting period, as expected from disinhibited MSNs, and observed through spiking activity (Fig 4I) and spectral content of population activity (Fig 4J). During PD with DBS under correlated noise, we found that this theta/beta firing of D1 and D2 MSNs is also preserved (theta frequency at 6.10 ± 0.62 Hz and beta frequency at 17.45 ± 1.09 Hz), as evidenced from the spiking activity (Fig 4J) and spectral content (Fig 4L). In our simulations, D1 MSNs and D2 MSNs fire synchronously. Indeed, the D1 MSNs and D2 MSN do not project to each other and receive FSI input with similar connectivity density. Adding mutual inhibition and asymetric FSI-projections will likely break the symmetry in MSN responses during the FSI off-cycle, bringing them closer to experimental findings on pro- and anti-kinetic firing properties as well as co-existent firing activity in other conditions [58]. We do not expect such a modification to alter the beta/theta activity pattern of the MSN populations, only the beta oscillation phase difference between the two MSN populations.

In the regime of synchronized noise, the DBS frequency also has an effect on the firing pattern of FSIs. We found that the FSI gamma firing also tracks the half-frequency of the DBS stimulation frequency (Fig 4M): the relation is linear within the range of 120Hz-150Hz, yet may break down outside this range. Furthermore, the theta frequency remains generally constant as DBS frequency changes (Fig 4N). More importantly, while DBS at low frequencies (e.g., around 65Hz) in uncorrelated noise conditions partially recovers dynamics observed in baseline condition (Fig S12C,D), it is unable to recreate in FSIs theta/gamma dynamics in correlated noise conditions (Figure S14A,B: example for DBS at 65Hz), and thereby does not recover beta bursts in MSNs (Figure S14C,D). Indeed, stimulation at low frequencies does not raise the FSI excitation enough to yield bursts of gamma, as observed via FSI membrane potentials (Fig S14E−H) or average firing rates (Fig S14I). FSI theta cycles are then not produced (Fig S14J). The effects of stimulation frequency on firing activity suggests that this parameter ought to be clinically adjusted to enable operation of natural FSI properties, and ensure as close as possible natural dynamics, which will be patient specific. A thorough study of these effects is left for another investigation.

## Discussion

Our studies suggest that a clinically beneficial effect of high-frequency DBS may be achieved through normalization of striatal dynamics via engaging the subthalmo-striatal pathway. Several effects emerge from stimulating this pathway that help restore functional striatal network dynamics including (i) reversal of D2 MSN over-excitability through increased FSI inhibition of MSNs, which in turn leads to (ii) decreased beta oscillations, (iii) decreased amplification of beta oscillations in the loop due to decreased excitability of the STN somas, an amplifier of the beta rhythm, and (iv) restoration of FSI theta/gamma network dynamics, normally seen in high-DA states in the non-parkinsonian striatum. Importantly, we find that the functionality of DA (patterning of MSNs to burst at beta only for the duration of a theta cycle) is restored and can be modulated in the absence of DA by the amount of correlation in the corticostriatal input. Thus, our studies attribute restoration of network dynamics in striatum to DBS engagement of the STN-to-striatum pathway.

As a summary, the effect of DBS in two-fold. DBS interrupts the amplification of beta activity around the loop by stopping STN somatic activity from reaching the FSIs. However, only interrupting the propagation of beta by suppressing STN is not enough to restore striatal function, as this cannot suppress the beta activity intrinsically generated by the striatum (Fig S5E,F: showing an absent STN-FSI connection). Stopping STN somatic activity from reaching the FSIs, however, re-positions the FSIs as potential striatal beta suppressors, and the axonal stimulation is then necessary to excite the FSIs and restore adequate inhibition onto the MSNs.

Computational work on the mechanisms of DBS in STN treatment has, thus far, focused on restoring normal functioning of the basal ganglia output, notably the GPi. In particular, work has shown that DBS can alter the input received by GPi to restore its functionality; this can be achieved by increasing regularity in GPe inputs which can reduce response variability in GPi neurons [6], or by the convergence of additional excitation from STN and inhibition from GPe that restore regular GPi firing [7], or more generally, through a mixture of responses caused by HFS converging in the GPi [59] regularizing GPi activity. It has also been proposed that DBS results in altered projection dynamics between BG nuclei, and then engage a mechanism of converging network-wide input onto the striatum, which then regularizes the activity in striatopallidal projections to the GPi to restore GPi function [12]. The focus on normalization of basal ganglia output is to increase thalamic relay reliability, seen as essential to restore normal functioning of the BG-thalamo-cortical loop [8–10]. The work [10] establishes how DBS can replace rhythmic inhibition (at tremor frequencies in the theta range) from GPi to the thalamus with regularized firing that, despite an increase in GPi inhibition frequency and amplitude, restores the responsiveness of thalamocortical cells to sensorimotor input. The work [11] builds on the model in [10] and offers a detailed study of the effect of stimulation frequency on thalamic relay reliability.

Our work focuses, instead, on restoring the dynamics of the BG nuclei, which then results to a restoration at the level of the BG output. Our reasoning is that while output nuclei and thalamic functioning are essential to normal function, the intricacies of the mechanisms of action selection and switching motor programs, and their interaction with DA that is disrupted during PD, rely on the richness of BG rythmic dynamics. Restoring striatal dynamics can then restore BG output function and thus reliability in thalamocortical relay.

### Normal striatal dynamics and their functional roles

Beta oscillations [14–17], gamma oscillations [18, 19] and theta oscillations [20, 21] are normally observed in a coordinated manner in striatal networks to drive behavior. Parkinsonian symptoms are associated with excessive beta oscillations and a loss of modulation of these rhythms impedes normal behavior. We show that DBS can restore these rhythms as observed in normal conditions, and can therefore aid in restoring and supporting the behavioral functionality associated to each.

Beta oscillations are a robust, task-modulated feature of healthy striatum in humans [60, 61], monkeys [14, 16, 17, 38, 39, 62], bats [63] and rodents [15]. They are also expressed in the membrane potential of MSNs in normal, non-parkinsonian rodents [64, 65] and MSN spike timing can be modulated in beta [17]. They tend to appear through bursts [38, 39]. The role of these bursts is thought to help switch among cell assemblies encoding motor programs [18, 41, 42]. In our model, MSN beta bursts in normal conditions emerge due to disinhibition during FSI quiescence periods following FSI gamma bursts. It is surges of dopamine that increase FSI excitation and electrical coupling, pushing FSIs to produce synchronized gamma bursts, followed by quiescence. If dopamine is sustained at high levels long enough, we observe that these gamma bursts are nested in theta cycles (Fig 4A).

Striatal gamma oscillations are considered to be generated through FSI activity [18, 66–69] and have been associated with the initiation and vigor of movement [70, 71]. Gamma activity has been found to have a positive correlation with dopamine and locomotion [37, 71–74] and to be anticorrelated in EEG and corticostriatal LFP to beta power [36, 71, 75]. Indeed, under high DA conditions, the gamma oscillations are nested within slow oscillations, leading to periodic gaps in FSI activity that allow MSN beta activity to emerge. We hypothesize that the FSI gamma bursts are necessary to terminate on-going motor programs, and initiate new programs by allowing for different MSN cell assemblies to activate, producing beta oscillations [22].

Theta oscillations are found in local field potentials recorded in the dorsal striatum [20, 76–78]. These rhythms show phase amplitude coupling with gamma activity [77, 79], are suggested to be orchestrated by FSI activity [20], and are modulated by dopaminergic activity and locomotion [20, 76]. These rhythms are modulated during task performance [80], and arise particularly during navigation tasks where they coexist with the hippocampal theta rhythms [81, 82]. The striatal theta is, however, considered to have a striatal origin [20] and its interaction with hippocampal rhythms can prove key in coordinating learning and memory [78, 83]. These striatal theta rhythms can prove key in controlling of voluntary behavior at a theta time scale relevant to sequencing of behaviors.

### Abnormal BG network dynamics in parkinsonian conditions

Our previous work [13] suggests that networks of medium spiny neurons (MSNs) in striatum are capable of producing beta oscillations under normal conditions which become exaggerated in the parkinsonian state. Specifically, the beta oscillations will be exaggerated under conditions of high cholinergic tone or under conditions that tonically increase MSN excitation, such as loss of dopaminergic stimulation of D2 receptors. Consistent with our modeling findings, increasing the striatal cholinergic tone of normal mice produced robust, exaggerated beta oscillation in both the striatal local field potential and in cortex [13, 84, 85] as well as parkinsonian-like behavioral deficits [84]. Exaggerated beta oscillations further emerge in the striatal LFP of parkinsonian rats [86, 87] and parkinsonian non-human primates [88, 89]. Particularly, indirect pathway MSNs have been found to synchronize their spiking at beta frequency in the dopamine-depleted striatum of rats [87]. While beta activity is amplified in PD, it is not constant, but is modulated in amplitude in the form of frequent beta bursts with longer durations [40, 90, 91]. This modulation, in our model during PD, can be achieved by variations in MSN excitability, through exogenous cortical or thalamic input (e.g., Fig S13A,B as an example).

Excessive beta oscillations extend beyond the striatum and are observed in several structures in the CBT loop [48] following dopamine depletion in rodent models of PD [50, 69, 76], in nonhuman primate models of PD [88, 92] and humans PD patients [93]. The indirect pathway is particularly implicated in these abnormal dynamics [87], and this implication highlights a special participation of MSNs in beta generation in rodents [87, 88] and in non-human primates [94]. While the literature had often reported increases in striatal firing rates connected to the increased beta activity [95], there has been recent contradictory results that report the absence of increased firing rates [96]. Our modeling suggests that an excessive beta oscillation in the striatum cannot be produced by additional MSN synchrony alone, without enhanced excitability and therefore firing rates. However, our results do not rely on an excessive increase, as strong beta oscillations can be observed with only a 4- or 5-fold increase in firing rates.

Moreover, it has been reported [97] that MSNs do not display beta oscillations in parkinsonian conditions following dopamine depletion, contradicting findings and conclusions of other works [76, 89]. However, [97] utilizes an MPTP intoxication procedures that leaves a parkinsonian nonhuman primate incapable of executing behavioral tasks. It is then uncertain whether or not MSN activity in such a state is altered compared to what is expected in regular parkinsonian conditions, where subjects are capable of tasks.

Abnormal beta activity is likely to simultaneously emerge from multiple sources in the BG, and the literature suggests that it can also arise from STN-GPe interaction [49, 98] and cortico-subthalamic patterning [49, 99, 100]. Our results are consistent with having multiple sources for beta activity, and we expect multiple sources to contribute to network dynamics. However, our work builds on an intrinsic source of beta activity residing in the interaction of the MSNs that, when unmodulated, hinders normal striatal dynamics. As such, if DBS can restore striatal dynamics it will necessitate a mechanism to act on that intrinsic source. As we found, simply halting beta activity propagated by STN is not enough to restore normal dynamics. Nevertheless, it has been argued that the striatum cannot produce the beta oscillation by itself because blockade of striatun to GPe does not eliminate the beta [101]. Instead, this study suggests that either the STN-GPe loop or the hyperdirect pathway are sources of beta in non-human primates. However, in [101], the blockade is confined to only a small region, targeting a small portion of the striatum to GPe projections, which we estimated from GPe size to be around 3%. This allows the rest of beta-producing striatal neurons to transmit beta downstream and engage the rest of the CBT loop, including STN, which our current study suggests can amplify beta throughout the indirect pathway loop via the STN-FSI connection pathway. In striking contrast to the above study, [102] suggests that either GPe or striatum or both may underlie beta generation in PD rodents, having ruled out both STN and cortex. Such contrasting conclusions suggest the methodologies used as well as their interpretations need to be carefully evaluated. Importantly, the interpretation of any such results will necessarily be dependent on an accurate representation of BG network pathways and cellular physiology. Our current study highlights that even a minor pathway (the STN-to-FSI pathway) may prove to be a key component of system-level dynamics.

### The mechanisms of DBS that restore normal dynamics

Dopaminergic modulation is lost during PD, disrupting the ability of FSIs to synchronize and provide gamma bursts. However, our results show that DBS can substitute the mechanism of dopaminergic control by that of cortical noise control. Once DBS reduces the beta activity that FSIs receive as input from the BG, cortical input becomes the major FSI provider of noise, and its correlation among FSIs dictates whether or not FSIs fire in synchrony. Importantly, we consider, in normal conditions, high cortical activity correlation to coincide with high DA levels, thus yielding an adequate substitution in PD.

This result is reminiscent of mathematical results on correlated noise-induced synchrony, where synchrony is achieved in coupled excitable systems (e.g., neurons coupled by gap junctions) at certain levels of correlated noise, but breaks at lower and higher intensity of noise [103–105]. Furthermore, regular input can break the synchrony created by correlated noise, suggesting an additional mechanims by which DBS can break synchronous beta oscillations in the BG loop [103].

While correlated noise can aid in sustaining synchronous theta/gamma firings among FSIs during DBS in PD, it can additionally increase beta synchrony at the level of FSIs during PD in the absence of DBS. An increase of beta synchrony has been suggested by [106], resulting from periodic correlated cortical input to FSIs. The beta sychrony in FSIs is considered in [106] to be sustained by the FSI gap junctions. In our work, we have not modeled any mechanism that can generate beta oscillation in FSIs from cortical connections; FSIs, in our work, are entrained at beta by STN projections. Regardless, although further weakening gap junctions might lessen beta oscillations in PD, it would be at the expense of a deficit in re-establishing the theta/gamma oscillations.

While such a cortical control provides a mechanistic fix in restoring theta/gamma FSI oscillations, relying only on cortical control might prove to problematic, as it might be unable to sustain the necessary activity for a long period of time to enable sequential tasks. This is particularly relevant in speech fluidity, in which we find verbal fluency to be impaired with DBS in STN [107, 108]. Additional theta stimulation, along with HFS, might prove to be an effective therapy to maintain the low oscillatory state.

DBS in STN will likely have direct effects on many connected and adjacent brain structures, with potential therapeutic effects. DBS in STN can excite the GPe, the GPi and the PPN [109] altering firing rates [34]. In these sites, stimulation has also proven to improve parkinsonian symptoms [110–113]. The GPe, in particular, has a normally high spiking rate, and STN stimulation may restore GPe functioning through a mechanism similar to what we observed for FSIs, by increasing firing rates and countering the effect of beta-oscillation on GPe through high frequency oscillations. The GPe projects back to the striatum, targetting MSNs and FSIs though both arkypallidal and prototypical cells[97, 114, 115], which may prove to be a complementary route to restore striatal dynamics. HFS has also shown to result in antidromic activity [116–118], notably at the level of cortex through the hyperdirect pathway.

Enforcing the beta generation to be striatal is not required for the efficacy of the proposed model of DBS. Our model of DBS acts by decoupling STN somatic activity from axonal activity, and replacing axonal activity by HFS [30]. It then stops the propagation of beta oscillations that are present in the STN, and increases the excitability of the FSIs to restore effective MSN inhibition. It blocks beta activity whether it is generated from the STN-GPe loop or exogenously introduced e.g., via cortical input [49]. The BG loop in particular acts as a beta resonator, and DBS in STN targets a central node in this loop that breaks beta resonance. STN itself can also be considered to be an amplifier of beta oscillations, and DBS i s capable of suppressing that amplification by decoupling axonal activity from somatic activity. Regardless, simply silencing STN is not enough to restore normal dynamics. It is essential to restore FSI function by adequate excitation, otherwise the MSNs will remain in their abnormal oscillatory state.

### Optimal parameters of DBS

The dependencies of STN DBS clinical efficacy on stimulation frequency, intensity and pulsewidth have been reproduced in our simulations, with optimal results appearing within the clinical range [119].

Gradually increasing stimulus intensity will result in oculomotor effects, autonomic effects, paresthesia, dystonic effects and speech impairment [54, 120–122]. These effects potentially emerge from a current spread to pyramidal tracts and other adjacent structures [123]. We do not study such pathological conditions, and limit the voltage to therapeutics ranges.

Pulse-widths have conventionally ranged between 60*μs* and 450*μs* for DBS in STN [119], though recent research has been investigating the effect of shorter pulse-width [124]. The choice of pulse-width additionally directly modulates the amount of excitation received by the FSIs, and dictates the amount of time that the AMPA gates are open. Our model suggests that an increase in pulse-width while keeping a fixed amount of excitation requires a decrease in stimulus intensity. This relation has been observed clinically while modifying pulse-width with the goal of keeping clinical efficacy unchanged [54].

Significant beneficial effects have been observed for stimulation frequencies ranging from 50Hz to 185Hz, with the most, observed and maximized, within the 130-185Hz range [54, 55]. Stimulation at low or beta frequencies have resulted in worsening of symptoms [54, 56, 57]. Consistent with this, our model suggests low-frequency DBS stimulation will lead to increased BG beta oscillations. We can expect optimal DBS clinical effectiveness when the stimulation frequency matches the natural oscillatory dynamics of FSIs, in the gamma range [18, 22, 125] and can resonate with FSI behavior. In the ranges above 120Hz, the FSIs oscillate at half of the stimulation frequency with an oscillation of 67.5Hz achieved for stimulation at 135Hz. Overall, we believe that choice of DBS frequency variations between patients may partly be due to individualistic differences in oscillatory gamma frequencies, which our model links to properties of the FSI D-current.

## Brief methods

All neurons are modeled using a single compartment with Hodgkin-Huxley-type dynamics. The voltage change in each cell is described by: 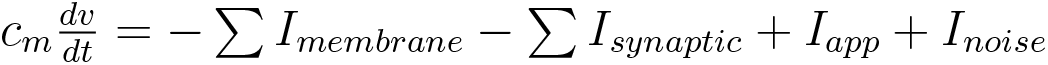. All cells display a fast sodium current (*I_Na_*), a fast potassium current (*I_k_*), a leak current (*I_L_*) for membrane currents (*I_membrane_*). MSNs additionally display an M-current and FSIs additionally display a D-current. The synaptic currents (*I_synaptic_*) depend on the connectivity. The aggregate population activity of MSNs and of FSIs, from which spectral information was determined, consisted of the sum of GABAa synaptic currents between MSNs and between FSIs, respectively. The aggregate population activity of STN and GPe consisted of the sum of membrane potentials of STN and GPe cells, respectively. Modeling details are provided in Supplementary Information. Our network models were programmed in C++ and compiled using GNU gcc. The differential equations were integrated using a fourth-order Runge-Kutta algorithm. The integration time step was 0.05ms. The model output was analyzed using Python 3.

## Acknowledgments

This work was partially supported by the NIH Award P01 GM118269 (to E.B. and N.K.).

## Author contributions

E.A., M.M., and N.K. designed the research; E.A. performed the research with input from E.B., N.K. and M.M.; E.A., E.B., N.K. and M.M. wrote the paper.

## A Supplementary Information

Below are the details directly referenced in the main document.

### A.1 Activity of isolated MSN network in baseline condition

In an isolated MSN network in baseline condition (Fig S1A), although there is no obvious synchrony in the MSN spiking activity (mean±SD of MSN firing rate: 1.46 ± 0.046 spk.s^−1^) (Fig S1B), the spectrum of population activity shows a peak in the low beta frequency range (mean±SD: 17.53 ± 2.77 Hz) (Fig S1C). Indeed, work from our lab [13] has shown that the MSN population can produce beta oscillations through an interaction between an inhibitory M-current and interneuronal GABAa inhibition.

### A.2 Activity in core striatal model in baseline conditions

Our simulations for the core striatal model (Fig S1D–F) in baseline conditions show sparse gamma oscillations in FSI activity, where cells do not fire at every gamma cycle but produce gamma oscillations as a population. The FSI firing is sparse and does not show periodicity in spiking activity at the level of a single cell (mean±SD of FSI firing rate: 10.66 ± 0.061 spk.s^−1^) (Fig S1E), but produces a peak at gamma frequencies (mean±SD: 58.63 ± 1.96 Hz) in the spectrum of population activity (Fig S1F). Work from our lab [22] has also shown that the FSIs are capable of intrinsically producing gamma oscillations due to their D-current, both at a population level and at the level of a single cell. We believe that this gamma oscillation generally plays a crucial role in suppressing beta oscillations in the striatum.

### A.3 Effect of the FSI-MSN projection

We find that adding FSI inhibition onto MSNs (Fig S1D) changes the MSN population activity spectrum from one showing a peak of low beta-band activity (Fig S1C) in the case of an isolated MSN population activity to one without beta-band activity (Fig S1F). The change in MSN spectral properties, after adding FSI inhibition, is not a result of a decrease in MSN firing rate, but a change in the dynamics driving MSNs to spike under FSI inhibition. The average MSN spiking rate instead slightly increases from mean±SD:1.46 ± 0.046 spk.s^−1^ without inhibition to 1.88 ± 0.057 spk.s^−1^ with FSI inhibition (Fig S1B,E). We believe that this increase is due to MSN rebound spiking from FSI inhibition: as we decreased FSI inhibition onto MSNs, by decreasing the maximal GABAa conductance of the FSI projections to MSNs, we first obtain a decrease in spiking activity (e.g., from 1.84 ± 0.06 spk.s^−1^ at *g* = 0.6 mS·cm^−2^ to 0.75 ± 0.05 spk.s^−1^ at *g* = 0.024 mS.cm^−2^) (Fig S2A). Decreasing the maximal conductance further eventually leads to an increase in MSN spiking activity (e.g., 0.98±0.05 spk.s^−1^ at *g* = 0.006 mS.cm^−2^) and a re-emergence of beta-band activity (Fig S2A), indicating an ineffective FSI inhibition at low conductances.

### A.4 Effect of adding GPe and STN in baseline conditions

Upon adding the GPe and STN populations, the spectra of population activity for MSNs and FSIs remain largely unchanged (comparing Fig S1F to Fig 1C). However, we observe that the MSN average firing rate decreased (p¡0.001, Welch’s t-test) from around 1.8 spk.s^−1^ (Fig S1E) to 1.21 ± 0.078 spk.s^−1^ (Fig 1B), and that the FSI spiking activity increased (p¡0.001, Welch’s t-test) from 10.66 spk.s^−1^ without STN input (Fig S1E) to 13.00 ± 0.067 spk.s^−1^ with STN input (Fig 1B).

The increase in FSI excitability and firing rate is followed by a slight increase (p¡0.001, Welch’s t-test) in FSI gamma frequency from 58.63Hz without STN input (Fig S1F) to 61.14±2.27Hz with STN input (Fig S1C), consistent with the correlation between FSI oscillatory frequency and cell excitability [22, 125]. The asynchrony of STN activity acts especially as a noise that further breaks any potential intrinsic FSI periodicity, thereby sustaining uncorrelated and irregular FSI firing. This irregularity, combined with a slight increase in FSI activity, leads to additional inhibition onto MSNs, lowering their spike rate.

### A.5 Activity of isolated MSN network and core striatal model in PD

As shown in previous work [13], in the presence of higher cholinergic modulation, an isolated MSN network (Fig S4A) is capable of producing strong beta oscillations, as evidenced directly by MSN spiking activity (Fig S4B) with an average spike rate of 5.32 ± 0.04 spk.s^−1^ and by the peak at the beta frequency 17.03± 0.47Hz in the spectrum of population activity (Fig S4C). Under parkinsonian conditions, in the presence of higher cholinergic modulation while providing additional D2 MSN excitability due to low levels of DA (Fig S5A), the spiking rate of MSNs is further increased to 7.66 ± 0.04 spk.s^−1^ (Fig S5B) while retaining a prominent peak at beta frequencies (at 20.02 ± 0.77Hz, higher than the frequency due to only an increase in cholinergic tone: p¡0.001, Welch’s t-test) in the spectrum of population activity (Fig S5C).

Similarly to our isolated MSN network, our core striatal model during PD (Fig S5D), comprised of MSNs and FSIs, is capable of producing a beta oscillation in MSNs, as weakly observed in the spiking activity (Fig S5E) and a peak at beta frequencies (18.77 ± 2.97Hz) in the spectrum of population activity (Fig S5F). However, the MSN beta activity is more spread out along the spectrum (Fig S5F) due to the presence of FSI inhibition. Although FSI activity is weakened in the PD case compared to baseline condition, as apparent from the sparse spiking activity (Fig S5E) and the spectrum of population activity (Fig S5F) where the gamma oscillation is no longer present, it is still capable of partially inhibiting the beta oscillation present in MSNs, causing changes in the spectral content of MSN firings, compared to that of an isolated MSN network. This change is also followed by a decrease in MSN firing rate from 7.66 spk.s^−1^ in an isolated MSN network (Fig S5B) to 6.62 ± 0.21 spk.s^−1^ in an MSN network with FSI inhibition (Fig S5E).

### A.6 Effect of adding GPe and STN in PD

After connecting the core striatal model to the STN and GPe, the beta frequency of the MSN activity (at 15.80Hz) decreased from that of the isolated MSN network (at 20.02Hz, p¡0.001, Welch’s t-test) and that of the core strial network (at 18.77Hz, p¡0.001, Welch’s t-test), suggesting that FSI activity is partially pacing MSN activity to fire at a lower frequency due to synchronized and increased FSI inhibition. This increase in beta band power in MSNs is also accompanied by a decrease in firing rates to 4.85 ± 0.13 spk.s^−1^ (p¡0.001, Welch’s t-test) (Fig 2B), a reduction likely due to the reduction in beta frequency despite the increase in beta activity in FSIs.

### A.7 Causes of the increase in MSN excitability

It is the increase of excitability of the MSNs that generates beta activity in the striatum (Fig S6A), resulting from a combined effect of a decrease in M-current conductance (*g_M_*) (Fig S6B), an increase in background excitation (*I_app_*) (Fig S6C) and a decrease in FSI GABAa inhibition (*g*_FSI-MSN_) (Fig S6D): separately restoring each of these changes decreases beta activity in the MSNs. The average MSN spiking rate decreases from 4.85 ± 0.16 spk.s^−1^ to 4.00 ± 0.10 spk.s^−1^ by restoring *g_M_* (p¡0.001, Welch’s t-test), to 4.14 ± 0.12 spk.s^−1^ by restoring *I_app_* (p¡0.001, Welch’s t-test) and to 4.73 ± 0.13 spk.s^−1^ by restoring *g*_FSI-MSN_ (p=0.0014¡0.005, Welch’s t-test) (Fig S6A–D).

### A.8 Resonance properties during PD

In PD, the MSN beta oscillation is synchronous enough to pattern the GPe at beta frequencies, as evidenced by the spiking activity (Fig 2B). The synchronous beta activity of GPe inhibits STN activity and, in turn, patterns it at beta frequencies, also evidenced by the spiking activity (Fig 2B). STN beta activity finally reaches the FSIs through the STN-FSI direct projections, and creates beta oscillations in the FSIs, weakly observed in spiking activity (Fig 2B) and more strongly in spectral properties (Fig 2C).

In baseline condition, the FSIs suppress, through their sparse gamma activity, the weak beta activity generated by the MSNs. However, if FSI activity is instead contaminated with beta activity, we expect the FSIs to become a conduit for beta-activity: the MSNs will resonate to the FSI beta-inhibition due to their intrinsic and network dynamics, and beta activity is then amplified in MSNs and throughout the loop. Particularly, in our simulations during PD, we find that the beta activity generated by MSNs is strong enough to reach STN through the GPe, as observed from spiking activity (Fig 2B) and population activity spectra (Fig 2C). The STN activity then patterns the FSIs at beta frequencies, and we observe resonance in the whole loop enabled by the FSIs: in the spectrum of MSN activity in the closed-loop system (Fig 2C), the power at the peak of beta-band activity (Fig 2C) is greater than that with STN input removed (p¡0.001, Welch’s t-test) (Fig S5F) and (ii) the modal beta oscillation frequency is lower than that of an isolated MSN network (p¡0.001, Welch’s t-test) (Fig S5C).

To further test the resonance property of the closed-loop system, we introduced direct exogenous beta activity into the MSNs during baseline condition. For high input amplitude, we observe resonance at the input frequency at the level of MSNs with the appearance of harmonics (Fig S7A). This resonance at high input amplitude appears in all four populations. As we decreased the input amplitude, the resonance at the level of MSNs weakens (Fig S7B) and ceases to exist (Fig S7C). More essentially, in baseline condition, we hypothesize that the closed-loop network has a capacity to dampen beta oscillations, because of the natural role of FSIs, instead of implementing positive feedback. To test this, we used a sustained high amplitude beta-activity as input into the MSNs for a predefined period and then removed that input. We find that the system returned from resonance to normal operation, in terms of spiking activity (Fig S8A) and spectral properties (Fig S8B).

### A.9 Effect of exogenous input to STN

We find that an additional exogenous input into STN, if provided at the resonating frequency (15.8Hz) amplifies the existing beta oscillations throughout the loop, increasing the average MSN firing rate to 5.83 ± 0.12 spk.s^−1^ (p¡0.001, Wlech’s t-test) (Fig S9A,B). When the beta input frequency is not the resonating frequency (e.g., 17.5Hz or 20Hz instead of 15.8Hz), the input entrains the BG oscillations, and they lock to the frequency and phase of the input (Fig S9C–D).

### A.10 Fixing stimlation intensity

We did not model the exact voltage value applied onto the axons, as it necessarily differs from the clinical parameter applied somatically. Instead, we assume that increasing the applied voltage at the level of the soma is equivalent to increasing the applied voltage at the level of the axon. The stimulation of the STN axons will induce AMPA currents in FSIs, via increases in the AMPA gating variables: the rate function for the open state of the AMPA receptor increases as the presynaptic axonal voltage increases, up to a maximum rate (See methods section below for more details). We thus fixed the applied voltage to the one that yields the maximum rate, and vary the remaining two parameters: stimulation frequency and pulse width.

### A.11 FSI gamma frequency during DBS

As we vary the stimulation frequency, we find that the FSIs produce bursts at half of the stimulation frequency (Fig 3D). This linearity breaks down as we increase (e.g., above 150Hz) the stimulation frequency (Fig 3D). Indeed, inspecting the spectrum of FSI population activity, outside the 120-150Hz range, reveals a less pronounced peak around the half-frequency (Fig S11A). We hypothesize that the bursting dynamics of the D-current impose a limit on the bursting frequency. As such, while 135Hz is too high a frequency for the FSI bursts to be entrained at, the FSIs naturally follow a bursting at around 67.5Hz, firing at most every other cycle.

### A.12 Effects of changing DBS frequency

DBS produces a sawtooth-shaped AMPA current at the stimulation frequency (Fig S11B). The stimulation frequency then dictates the period between the peaks of AMPA currents, but also changes the baseline of added excitation: the higher the frequency, the less time the current has to decay (Fig S11B). This current leads to fluctuations in the membrane potential of the FSIs (Fig S11C), and has an effect of increasing the overall excitability of the FSIs, as observed through an increase in FSI firing rate (Fig 3E). The increased firing rate of FSIs increases the amount of inhibition on MSNs, leading to a decrease in firing rate as the stimulation frequency is increased (Fig 3F).

### A.13 Effects of changing DBS pulse width

Changing the pulse width, while fixing the stimulation frequency, modifies the duration in which the AMPA current rises before it decays again (Fig S11D), and thus modifies how much of an effect DBS stimulation has on FSI activity. Decreasing the pulse width decreases the level of excitation onto FSIs as observed through a decrease in firing rate (Fig S11E), leaving its inhibition onto MSNs ineffective for low values and allowing beta-activity to re-emerge. Indeed, we observe an increase in MSN average firing rate (Fig S11F) as the pulse width is decreased.

### A.14 Effects of STN input on FSIs

The loss of DA during PD renders the FSIs unable to achieve a state similar to that of high DA levels during normal condition. This is due to two reasons. First, increasing the background excitation onto FSIs drives them at a theta/gamma at the level of a single cell. In PD, the FSIs lose the potential excitation received from high DA levels to change their firing pattern. Second, whether or not FSIs are able to synchronize is governed by how strong the gap junctions are with respect to external noise that the FSIs receive. The loss of DA severely reduces the electrical conductance of the gap junction [52] and compromises the ability of FSIs to synchronize at theta/gamma, especially in the presence of beta oscillations. DBS acts at these two levels, and effectively restores FSI synchrony by combining two effects. First, through HFS, DBS increases the level of FSI excitation to drive them at a theta/gamma oscillation at the level of a single cell. Indeed, we can observe gamma bursts in spiking activity (Fig 3B), and peaks at theta/gamma frequencies in the spectral properties of population activities (Fig 3C). Second, by additionally disconnecting the FSIs from STN, DBS reduces the amount of noise that reaches the FSIs from STN, allowing the weakened gap junctions to then synchronize FSI theta/gamma activity.

Specifically, STN provides noisy input to the FSIs, through its direct projections. In normal condition, the STN firing is tonic and asynchronous, providing the FSIs with noisy input that is uncorrelated among FSIs. However, the gap junctions of the FSIs, in normal conditions, are strong enough to overcome this noise, and achieve FSI synchrony at high DA levels. In parkinsonian conditions, STN is inputing beta activity into the FSIs, acting as noise with respect to desired synchronized theta/gamma oscillations. The electrical conductance of the gap junction, already weakened, leave the FSIs with little ability to synchronize in the presence of these beta oscillations. During PD, DBS dissociates STN axonal activity from STN somatic activity. The level of FSI synchrony is then controlled by how much the remaining background noise received by FSIs (which we modeled as cortical input) is uncorrelated/correlated among them.

### A.15 Failure of FSIs to yield theta/gamma oscillations in PD without DBS

When we provide FSI-correlated noise, FSI excitation is too low to achieve theta/gamma oscillations at the level of a single cell as observed in spiking activity (Fig 4E), and FSI activity is strongly contaminated with beta activity instead as observed in spiking activity (Fig 4E) and spectral content (Fig 4F). We also cannot achieve synchronous firing (Fig 4G) under FSI-correlated noise, even if we artificially increase excitation to have each FSI at a theta/gamma individually, as evidenced through the spectrum (Fig 4H).

## B Methods

### Model of neuron and dynamics

All neurons are modeled using a single compartment with Hodgkin-Huxley-type dynamics. The voltage change in each cell is described by:

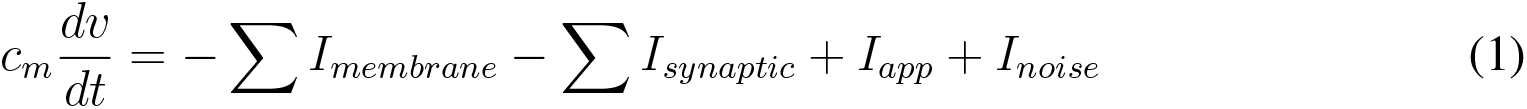

The membrane capacitance (*c_m_*) is normalized to 1 *μ*F.*cm*^−2^ for all neurons. All cells have a fast sodium current (*I_Na_*), a fast potassium current (*I_k_*), a leak current (*I_L_*) for membrane currents (*I_membrane_*). MSNs additionally have an M-current and FSIs additionally have a D-current (discussed in detail later). The synaptic currents (*I_synaptic_*) depend on the connectivity, and are discussed in the section on network connectivity and synaptic currents. The applied current (*I_app_*) is a constant that represents background excitation and the noise current (*I_noise_*) corresponds to a gaussian noise.

### Membrane currents and background excitation in baseline condition

The membrane currents are modeled using Hodgkin-Huxley-type conductance dynamics and formulated as:

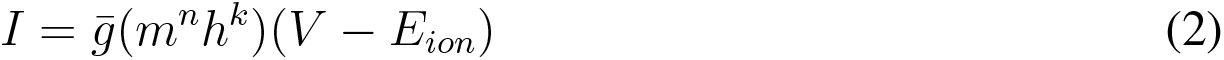

Every membrane current has a constant maximal conductance 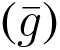 and a constant reversal potential (*E_ion_*). The activation (*m*) and inactivation (*h*) gating variables have *n^th^* and *k^th^* order kinetics with *n, k* ≥ 0. The dynamics of each gating variable evolves according to the kinetic equation (written here for the gating variable *m*):

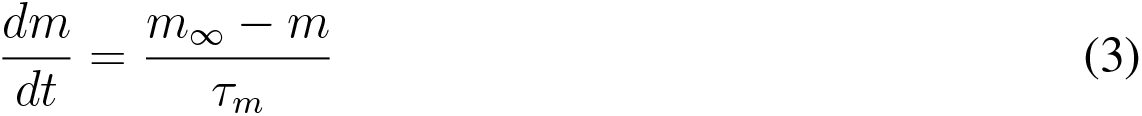

The steady-state function (*m*_∞_) and the time constant of decay (*τ_m_*) can be formulated as rate functions for each opening (*α_m_*) and closing (*β_m_*) of the ionic channel by using:

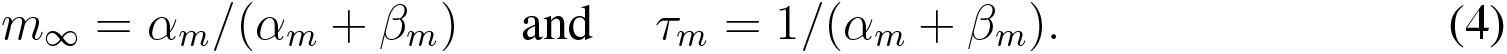

The baseline excitation (modulated by dopamine and acetylcholine) and the sum of all excitatory and inhibitory exogenous inputs for a given neuron (e.g., from the cortex, thalamus and non-modeled input) is introduced into the model using a constant background excitation term (*I_app_*). To account for variability in background excitation, we further introduce a Gaussian noise term (*I_noise_*). The Gaussian noise has mean zero and standard deviation dependent on the neuronal cell type.

#### Striatal medium spiny neurons (MSN)

Our model striatum consists of a network of medium spiny neurons (MSNs), which comprise about 95% of the neurons in the rodent striatum [126]. The membrane currents (*I_membrane_*) of MSNs consist of a fast sodium current (*I_Na_*), a fast potassium current (*I_K_*), a leak current (*I_L_*), and an M-current (*I_m_*) [127]. We do not model MSN up and down states which are not prevalent in the awake state [128], the state being modeled, and therefore we do not include the Kir current in our model, which is active during the MSN down state.

##### Fast sodium current

The sodium current (*I_Na_*) has three activation gates (n=3) and only one inactivation gate (k=1). The rate functions for the sodium current activation (*m*) and inactivation (*h*) variables are formulated as:

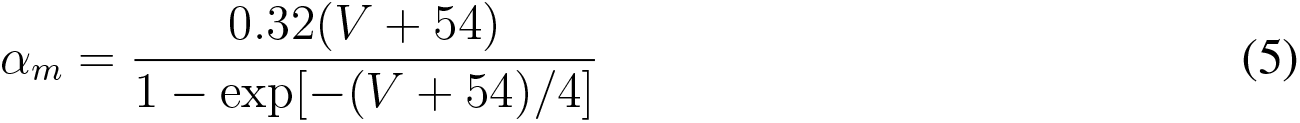

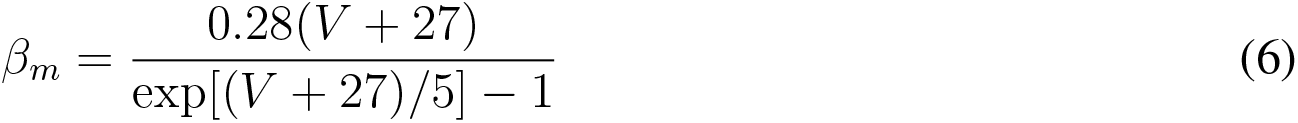

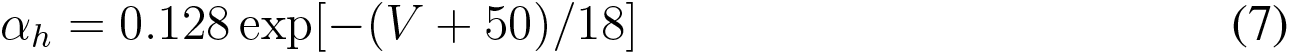

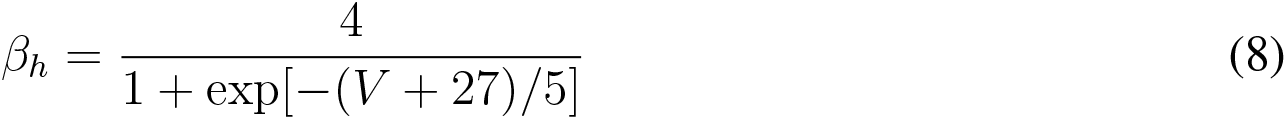

The maximal conductance of the sodium current is 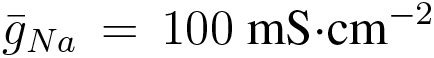. The sodium reversal potential is *E_Na_* = 50mV.

##### Fast potassium current

The fast potassium current (*I_K_*) has four activation gates (*n* = 4) and no inactivation gates (*k* = 0). The rate functions of the activation gate are described by:

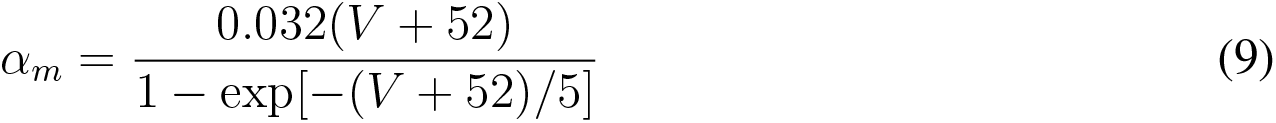

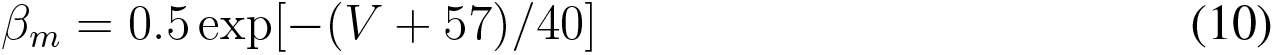

The maximal fast potassium channel conductance is 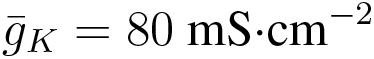. The reversal potential for potassium is *E_K_* = −100mV.

##### Leak current

The leak current (*I_L_*) has no gating variables (*n* = 0, *k* = 0). The maximal conductance of the leak channel is 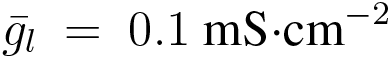. The leak channel reversal potential is *E_L_* = −67mV.

##### M-current

The M-current (*I_M_*) has one activation gate (*n* = 1) and no inactivation gate (*k* = 0). The rate functions for the M-current activation gate are described by:

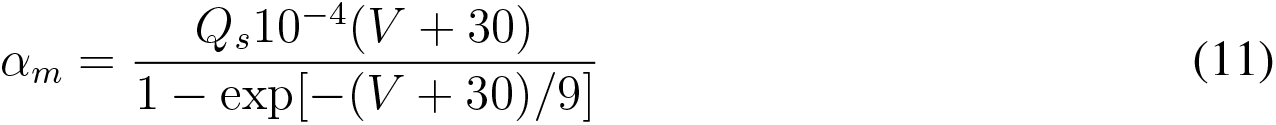

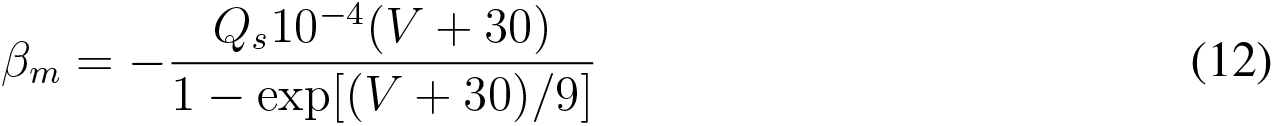

We use a *Q*_10_ factor of 2.3 to scale the rate functions of the M-current since the original formulation of these kinetics described dynamics at 23°*C* [129]. Thus, for a normal body temperature of 37°*C*, the M-current rate equations are scaled by *Q_s_*, which is formulated as:

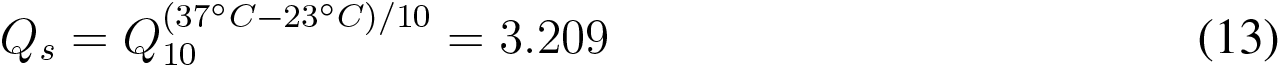

The maximal M-current conductance is 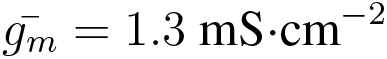 in baseline conditions.

##### Applied current and noise

The applied current (*I_app_*) is set to 1.19 *μ*A.*cm*^−2^, and the Gaussian noise (*I_noise_*) has mean 0 and standard deviation 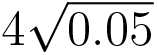 where 0.05ms corresponds to the time step of integration in our simulations.

#### Striatal fast spiking interneurons (FSI)

Striatal fast spiking interneurons (FSIs) were modeled as in [125] and [22], using one compartment. The membrane currents (*I_membrane_*) of FSIs consist of a fast sodium current (*I_Na_*), a fast potassium current (*I_K_*), a leak current (*I_L_*), and a D-current (*I_D_*). The formulation of these currents is taken from [22] that build upon previous models of striatal FSIs [125, 130].

##### Fast sodium current

The sodium current (*I_Na_*) has three activation gates (n=3) and only one inactivation gate (k=1). The steady state functions for the sodium current activation (*m*) and inactivation (*h*) variables and their time constants are described by:

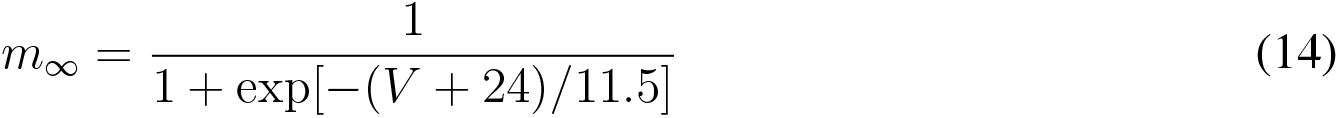

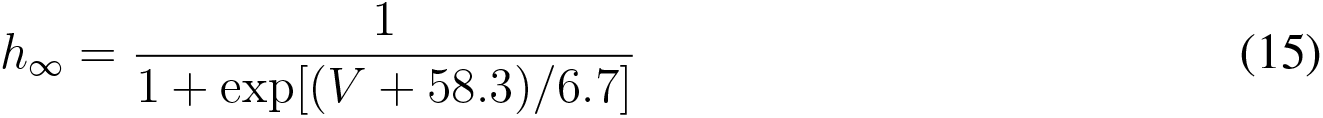

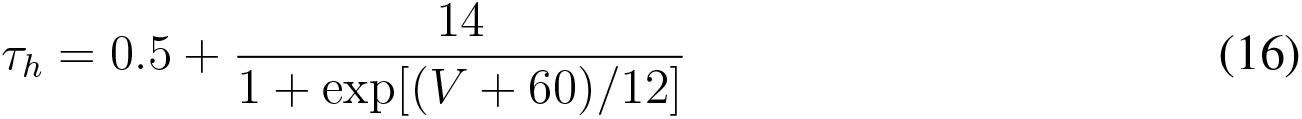

The time constant *τ_m_* is assumed to be negligible (around 0ms) and thus we set *m = m*_∞_ [125]. The maximal conductance of the sodium current is 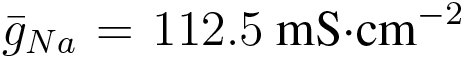. The sodium reversal potential is *E_Na_* = 50mV.

##### Fast potassium current

The potassium current (*I_K_*) has two activation gates (n=2) and no inactivation gate (k=0). The steady state function for the potassium current activation (*n*) and its constant (*τ_n_*) are described by:

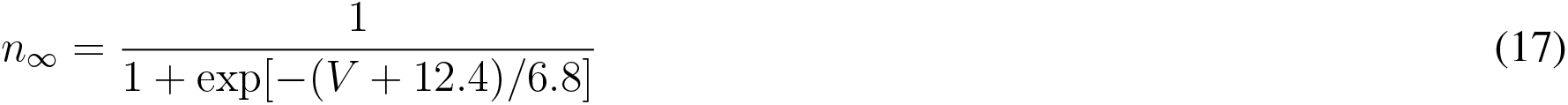

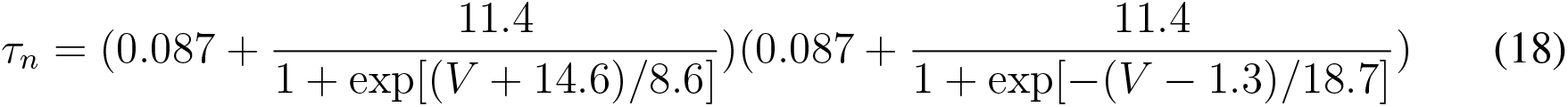

The maximal conductance of the sodium current is 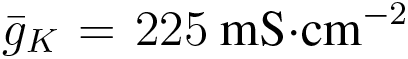. The potassium reversal potential is *E_Na_* = −90mV.

##### Leak current

The leak current (*I_L_*) has no gating variables (*n* = 0, *k* = 0). The maximal conductance of the leak channel is 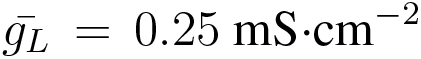. The leak channel reversal potential is *E_L_* = −70*mV*.

##### D-Current

The fast-activating, slowly inactivating potassium D-current (*I_D_*) is described mathematically as in [125] and has three activation gates (n = 3) and one inactivation (k = 1) gate. The steady state functions for the activation (m) and inactivation (h) variables and their time constants (*τ_m_* and *τ_h_*, respectively) are described by:

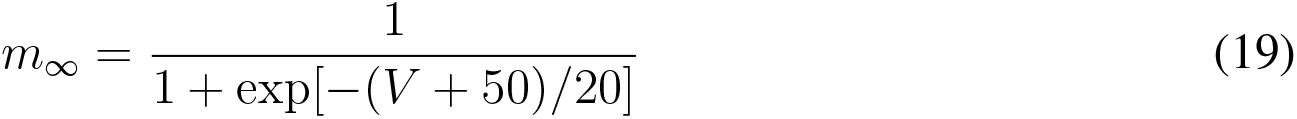

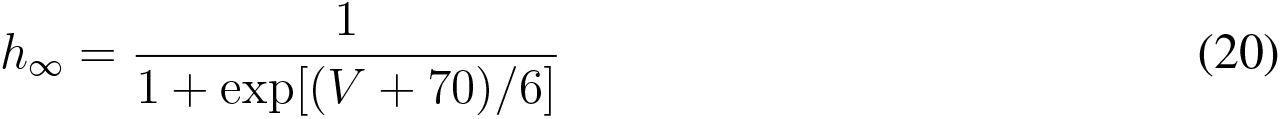

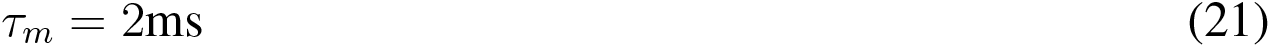

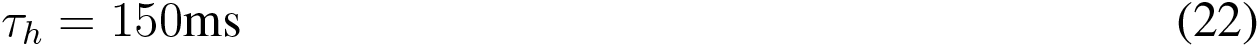

The maximal conductance of the D-current is 6 mS.cm^−2^.

##### Applied current and noise

The applied current (*I_app_*) is set to 5.5 *μ*A *cm*^−2^, and the Gaussian noise (*I_noise_*) has mean 0 and standard deviation 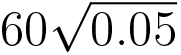 where 0.05ms corresponds to the time step of integration in our simulations.

#### Subthalamic and Pallidal neurons (STN and GPe)

The membrane currents for the subthalamic neurons (STN) and pallidal neurons (GPe) were modeled in a minimal manner, with just enough complexity to act as relay cells. The membrane currents (*I_membrane_*) of STN and GPe neurons are restricted to consist of a fast sodium current (*I_Na_*), a fast potassium current (*I_K_*) and a leak current (*I_L_*). We set the dynamics and parameters of these currents to be the same as those of MSNs, considered standard. Changes to the parameters will alter spiking waveform and neuronal excitability of these populations, and will thereby not affect our results as these are focused on MSN and FSI dynamics.

##### Applied current and noise

The applied current (*I_app_*) is set to 1.9 *μ*A.*cm*^−2^ for STN and 3.0 *μ*A.*cm*^−2^ for GPe, and the Gaussian noise (*I_noise_*) has mean 0 and standard deviation 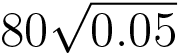) for both STN and GPe where 0.05ms corresponds to the time step of integration in our simulations. These values were chosen to give the GPe and STN populations appropriate baseline spike rates (~80Hz for GPe and ~ 17Hz for STN) and to mimic the largely asynchronous spiking seen under baseline conditions in these populations.

#### Network connectivity and synaptic currents in baseline condition

Our network consists of 100 D2 MSNs, 50 FSIs, 40 STN cells and 80 GPe cells. We have a total of 6 types of projections between the populations, 5 inhibitory (MSN-MSN, FSI-MSN, FSI-FSI, MSN-GPe and GPe-STN) and 1 excitatory (STN-FSI). All inhibitory synapses were modeled using GABAa currents and all excitatory connections were modeled using AMPA currents. We modeled the GABAa current (*I_GABAa_*) using a Hodgkin-Huxley-type conductance:

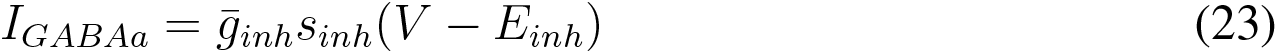

of the gating variables from all pre-synaptic connections. The gating variable *s_inh_* for inhibitory GABAa synaptic transmission is the sum:

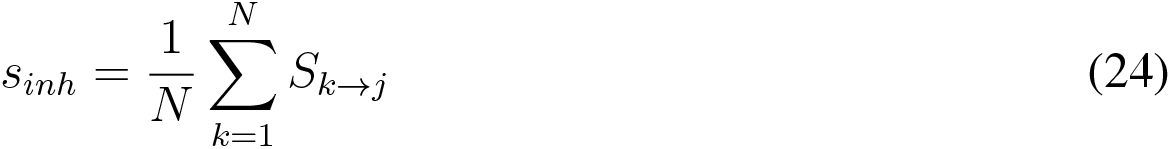

where *N* is the number of presynaptic neurons and *S_k→j_* describes the kinetics of the gating variable, for each pair of presynaptic neuron *k* and postsynaptic neuron *j*, evolving according to:

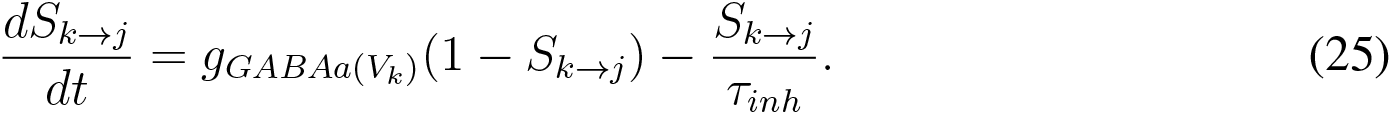

**Table S1:** Parameters for membrane currents and background excitation in baseline condition for all four neuronal populations.

|  | MSN | FSI | STN | GPe |
| --- | --- | --- | --- | --- |
| $C_m(\mu F \cdot cm^{-2})$ | 1 | 1 | 1 | 1 |
| $I_{app}(\mu A \cdot cm^{-2})$ | 1.19 | 6.2 | 1.9 | 3 |
| $g_{Na}(mS \cdot cm^{-2})$ | 100 | 112.5 | 100 | 100 |
| $g_K(mS \cdot cm^{-2})$ | 80 | 225 | 80 | 80 |
| $g_L(mS \cdot cm^{-2})$ | 0.1 | 0.25 | 0.1 | 0.1 |
| $g_M(mS \cdot cm^{-2})$ | 1.3 | | | |
| $g_D(mS \cdot cm^{-2})$ | | 6 | | |
| $g_{elec}(mS \cdot cm^{-2})$ | | 0.15 | | |
| $E_{Na}(mV)$ | 50 | 50 | 50 | 50 |
| $E_K(mV)$ | -100 | -90 | -100 | -100 |
| $E_L(mV)$ | -67 | -70 | -67 | -67 |

Note that *S_k→j_* is a function of the presynaptic voltage *V_k_*, and its dynamics depend on the dynamics of the presynaptic neuron *k*. The rate functions for the open state of the GABAa receptor (*g_GABAa_*(*V_k_*)) is described by:

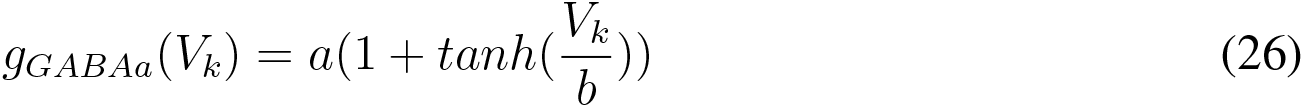

for values of *a* and *b* that depend on the pre- and post-synaptic neuronal cell types and are provided in Table S2. The maximal conductance 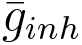 is scaled by the number of pre-synaptic neurons of a specific cell type. Excitatory AMPA synaptic currents use the same set of equations as for the GABAa current (eqns. 23 – 26) with the “GABAa” subscript replaced by “AMPA” and the “inh” subscript replaced by “exc”. Parameters for all synaptic currents are provided in Table S2.

##### Remark regarding D1 and D2 MSNs

We do not include the D1 MSNs in the context of PD, since they are believed to be much less active in this diseased state and are canonically considered to not be part of our modeled loop going through the indirect pathway of the BG. We only introduce D1 MSNs to establish how DBS in STN restores striatal theta/gamma oscillations and beta bursts during PD. When D1 MSNs (N=100) are introduced, they received the same projection pattern as MSN-MSN projections and FSI-MSN projections. In this work, we assume that the D1 MSNs do not project to GPe and thereby do not alter BG loop dynamics.

**Table S2:** Connectivity parameters in baseline condition. **\*** We specified the conductances for the MSN-GPe and GPe-STN projection such that (i) the MSNs have a capability of inhibiting GPe and GPe neurons have a capability of inhibiting STN, and (ii) the firing rates are preserved, as found in the literature, for STN and GPe during the various conditions. The tonic activity in STN and GPe is driven by a high applied current in our model, and the conductances have to be adapted, within bounds as observed in the literature, to ensure that synaptics currents can overcome the set high applied excitation. The exact choice of the conductance does not influence our results, as long as (i) and (ii) are satisfied. **\*\*** This projection is understudied in the literature. We estimated the parameters from the findings in [28].

| | $\bar{g}_{inh}$ | $\tau_{inh}$ | $E_{inh}$ | $g_{GABAA}(V_k)$ | Fraction | References |
| --- | --- | --- | --- | --- | --- | --- |
| MSN-MSN | 0.1 | 13 | -80 | $2(1 + \tanh(V_k/4))$ | 30% of 100 | [13, 131] |
| FSI-MSN | 0.6 | 11 | -80 | $4(1 + \tanh(V_k/10))$ | 15% of 50 | [22, 132, 133] |
| FSI-FSI | 0.6 | 6.5 | -80 | $4(1 + \tanh(V_k/10))$ | 58% of 50 | [22, 133] |
| MSN-GPe | 2.5* | 13 | -80 | $2(1 + \tanh(V_k/4))$ | 33% of 100 | [115] |
| GPe-STN | 0.3* | 10 | -80 | $2(1 + \tanh(V_k/4))$ | 5% of 80 | [134, 135] |
| | $\bar{g}_{exc}$ | $\tau_{exc}$ | $E_{exc}$ | $g_{AMPA}(V_k)$ | Fraction | References |
| STN-FSI** | 0.165 | 2 | 0 | $5(1 + \tanh(V_k/4))$ | 10% of 40 | [28, 132, 134] |

##### Remark regarding the FSI-MSN projections

Each of the 100 D2 MSNs randomly receives projections from 15% of the full FSI population (N=50). Experimental findings show that MSN receives projections from local FSIs (within a 250*μm* radius) with probability 45% [133]. We lower the connection probability to account for the ratio of MSNs:FSIs in our model network which is smaller than that found in vivo. While we can reduce the number of FSIs, we risk of losing heterogeneity in FSI responses. Instead, we will consider only a third of the FSIs to be local to an MSN and enforce the MSN to receive a projection from 45% of that third (which is equivalent to 15% of the FSI population). This will bring the number closer to experimental findings while keeping enough heterogeneity in responses.

##### Remark regarding FSI-FSI connectivity

FSIs were additionally connected by electrical connections via gap junctions. The electrical coupling for neuron j, has units in *μ*A.*cm*^−2^ and is formulated as:

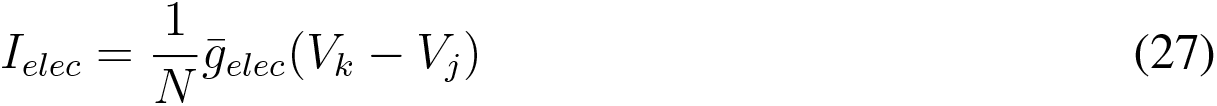

with *N* equal to the number of FSIs neuron *j* is coupled to. This yields a change:

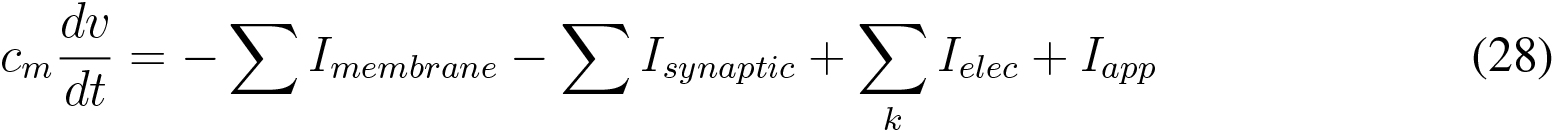

The value of the gap junction conductance 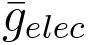 depends on the level of dopamine, and is set in baseline condition to 0.15 mS.cm^−2^. Each FSI was randomly electrically coupled to 33% of the remaining FSIs.

## Perturbations in Parkisonian conditions

Parkinson’s disease was modeled starting from baseline condition and adding the following biophysical changes. The applied current (*I_app_*) was increased to 1.25 *μ*A.*cm*^−2^ for MSNs [13] and decreased to 4.3 *μ*A.*cm*^−2^ for FSIs [22]. The maximal conductance for the MSN M-current was decreased to 1.2 mS.cm^−2^ [13]. The maximal conductance of the FSI-MSN projection was decrease by 20% to 0.48 mS.cm^−2^, as a result of the increase in cholinergic tone[53]. The maximal conductance of the FSI-FSI projection was also increased to 0.2. The value of the FSI electrical conductance *g_elec_* was also decreased to 0.075. The changes were extrapolated from [22], based on the changes in FSI parameters as a function of dopamine level.

When modeled, D1 MSNs inherit the same parameters as D2 MSNs except for *I_app_* which is decreased to 1.13 in Parkinsonian conditions.

## Applying DBS in Parkisonian conditions

Applying DBS during parkinsonian condition was modeled by applying a high frequency voltage input, directly altering the rate function *g_AMP A_*(*V_k_*) for the open state of the AMPA receptor governing the AMPA synaptic current of FSIs coming from STN. In particular we have:

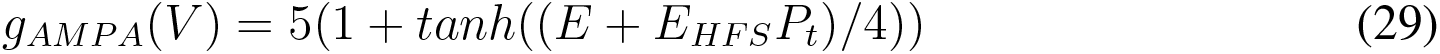

where *E* is set to −67*mV* that correspond to the resting potential of the neuron, *E_HF S_* is a constant voltage amplitude and *P_t_* corresponds to periodic pulses (modeled by a rectangular wave function of unit amplitude) with specified pulse-frequency (DBS frequency) and pulse-width (DBS pulsewidth, dictating the duty cycle). For all voltages *V*, we have −1 ≤ *tanh*(*V*) ≤ 1. The maximal *E_HFS_* was chosen so that *g_AMPA_*(*V*) ≃ 0 as *tanh*(*E/*4) ≃ −1 during the off-cyle of the rectangular wave *P_t_*, and *g_AMP A_*(*V*) ≃ 10 as *tanh*((*E + E_HFS_*)*/*4) ≃ 1 during the on-cycle of the rectangular wave. Therefore, different choices for the value of E do not alter the behavior of the model as long as they are sufficiently negative.

## High dopamine state and cortical input

A high dopamine state during normal condition was modeled starting with the parameters in baseline condition and adding the following biophysical changes. The applied current (*I_app_*) was decreased to 1.13 *μ*A.*cm*^−2^ for MSNs (expressing the D2-receptor) and increased to 8 *μ*A.*cm*^−2^ for FSIs. The maximal conductance of the FSI-FSI projection was also decreased to 0.05 mS.cm^−2^. The value of the FSI electrical conductance *g_elec_* was also increased to 0.3 mS.cm^−2^.

A high dopamine state is coupled with increased correlated cortical activity that leads to correlated noise onto FSIs. The noise term (*I_noise_*) for FSI neuron *j* is formulated as:

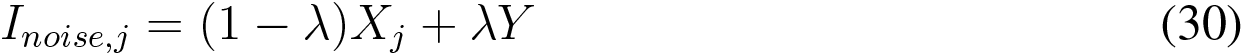

where *X_j_* is Gaussian random variable with mean 0 and standard deviation 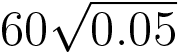 that is neuron specific and independent from that of <u>the</u> other neurons, and *Y* is Gaussian random variable with mean 0.4 and standard deviation 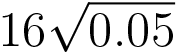 that is common to all FSIs. The parameter *λ* is fixed between 0 and 1 and dictates the amount of correlation in the FSI noise. We modeled the baseline condition with only uncorrelated noise with a *λ* = 0. We modeled the normal condition with high dopamine levels to consist primarily of correlated noise with a *λ* = 0.9.

D1 MSNs are modeled in the same manner as D2 MSN but with *I_app_* increased to 1.23 *μ*A.*cm*^−2^ in normal conditions with high dopamine levels, and decreased to 1.13 in Parkinsonian condition.

## Population activity and local field potentials

Oscillations have been typically detected in local field potential (LFP) measurements reflecting aggregate population activity. We sought similar signals and dissected their spectral content. A striatal LFP will necessarily contain a mixture of MSN and FSI activity, with MSN activity accounting for most of the signal given the size of the FSI population compared to the MSNs. To assess the activity of the separate populations, we thus defined an aggregate signal for each.

Synaptic currents have been used in models of LFP, EEG, or MEG [136–138]. Therefore, per [13], we model the aggregate population activity of MSN as the sum of all GABAa currents between MSNs. The MSN GABAa currents in our model reflect the spiking activity of MSNs, since each time an MSN spikes, it produces a GABAa current in other MSNs. Thus, our aggregate signal is tracking the population spiking activity of the MSNs. We are particularly interested in the population spiking activity of MSNs since this is the signal that will be transmitted to the output structure of the striatum, namely the GPe.

The FSIs also inhibit one another, and our aggregate signal consists of the GABAa synaptic currents between the FSIs. That aggregate signal should reflect the same spectral content as the sum of GABAa synaptic currents arising from the projections from FSIs to MSNs.

We have not modeled any connections between GPe or STN cells, and as such, the aggregate signals for GPe and STN were taken to be the sum of the membrane potentials of GPe and STN neurons, respectively.

To calculate the power spectral density of the LFP signal, we derive the discrete Fourier transform of the aggregate signal computed via the fast fourier transform implemented in Python SciPy. Power spectral density plots were constructed from simulations run for 5.5 seconds, while discarding the first 200ms of a simulation to ensure that the activity stabilized from the initial conditions.

## Statistics

Plots of spiking activity and spectral content in all conditions were drawn from 5.5 second simulations. Statistics and statistical test were computed using Python 3. The results (e.g., averages and standard deviations) in each condition were based on 25 simulations. Parametric sweeps were based on 10 simulations per parameter value, unless otherwise indicted.

## Simulation methods

Our network models were programmed in C++ and compiled using GNU g++. The differential equations were integrated using a fourth-order Runge Kutta algorithm. The integration time step was 0.05 ms. Model output is graphed and analyzed using Python 3.

## C Supplementary Figures

**Figure S1:**
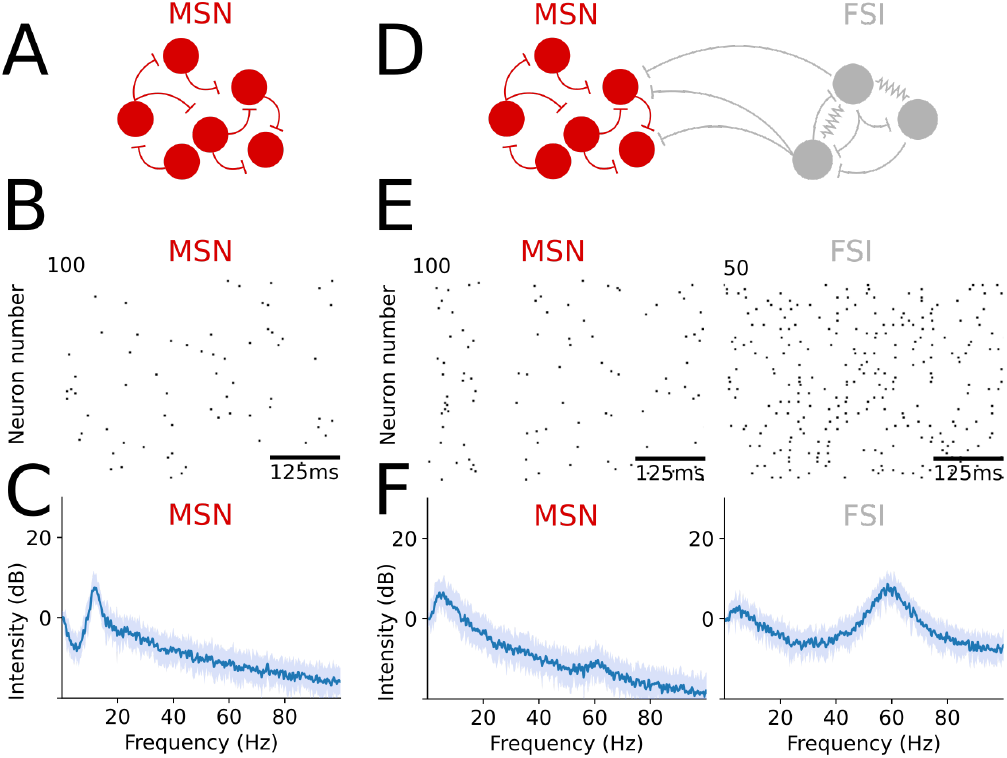
Population dynamics in baseline condition, in the core striatal model. **(A)** Schematic illustrating the MSN network, unconnected to other populations. **(B)** Raster plot showing spiking activity of MSNs in baseline condition for the network in (A) (MSN firing rate: mean SD=1.46 0.046 spk.s^−1^, N=25 simulations). **(C)** Graph showing the average (blue) and standard deviation (light blue) of the spectrum of MSN population activity, all in baseline condition for the network in (A) (N=25 simulations). **(D)** Schematic illustrating the core striatal model. **(E)** Raster plots showing spiking activity of MSN and FSI neurons, all in baseline condition for the network in (D). MSN firing rate: mean SD=1.88 0.057 spk.s^−1^, N=25 simulations. FSI firing rate: mean SD=10.66 0.061 spk.s^−1^, N=25 simulations. **(F)** Graph showing the average (blue) and standard deviation (light blue) of the spectra of MSN and FSI population activity, all in baseline condition for the network in (D) (N=25 simulations).

**Figure S2:**
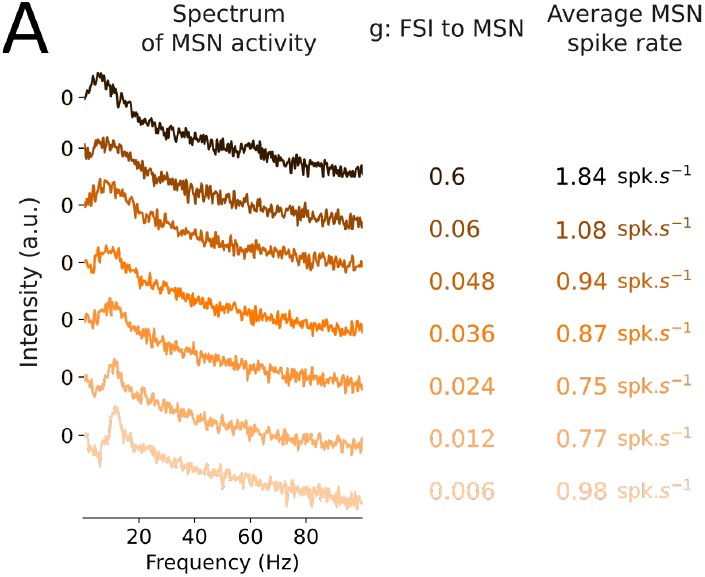
Changes in MSN activity as a function of FSI inhibition. **(A)** Graphs showing the spectrum of MSN population activity and their average firing rate in the core striatal model during baseline condition with different maximal GABAa conductances for the FSI projections to MSNs (N=10 simulations for each value of maximal GABAa conductance.

**Figure S3:**
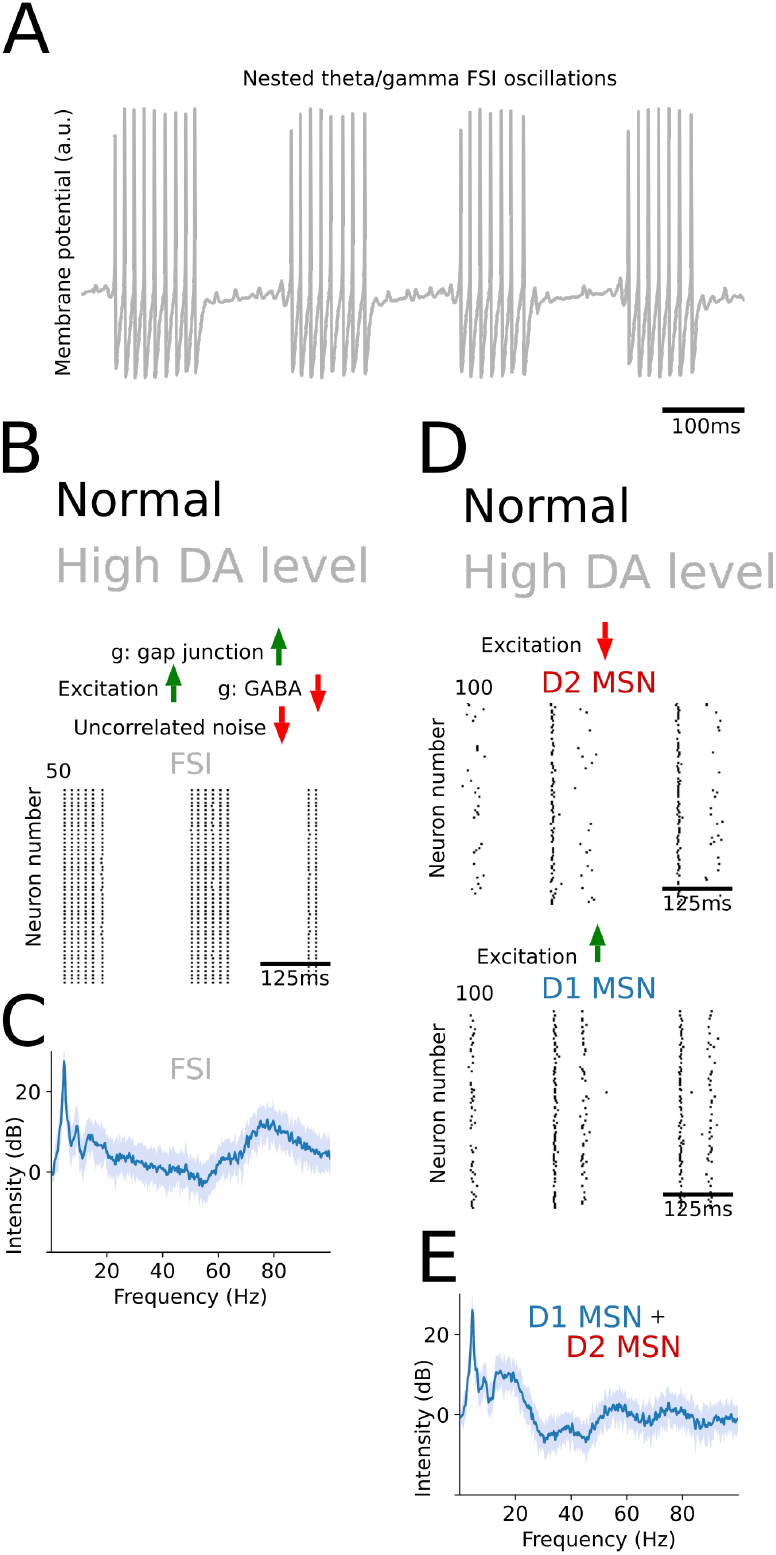
Population dynamics for high levels of dopamine in the core striatal model. **(A)** Graph showing theta/gamma oscillations of an isolated FSI subjected to high excitation due to an increase of dopaminergic level in the Striatum. **(B)** Raster plot showing spiking activity of FSI neurons, in normal conditions with high level of DA, in the core striatal model comprising only FSIs and MSNs. **(C)** Graph showing the average (blue) and standard deviation (light blue) of the spectrum of FSI population activity, in normal conditions with high level of dopamine, in the core striatal model comprising only FSIs and MSNs. **(D)**, **(E)** Graphs displaying population activity of MSN neurons, as in (B) and (C), for normal conditions with high level of dopamine, in the core striatal model. The raster plots show D1 and D2 MSN activity separately.

**Figure S4:**
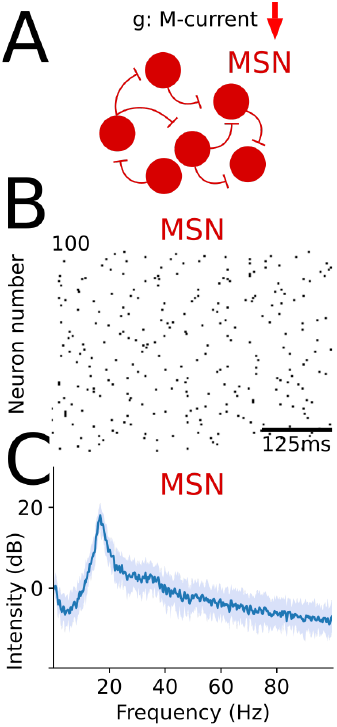
Population dynamics of an MSN network during increase in cholinergic tone. **(A)** Schematic illustrating the parametric changes in the MSN network during high cholinergic tone from parameters in baseline condition. **(B)** Raster plot showing spiking activity of MSNs during high cholinergic tone condition for the network in (A) (MSN firing rate: mean SD=5.32 0.037 spk.s^−1^, N=25 simulations). **(C)** Graphs showing the average (blue) and standard deviation (light blue) of the spectrum the MSN population activity, during Parkinsonianhigh cholinergic tone condition for the network in (A) (N=25 simulations).

**Figure S5:**
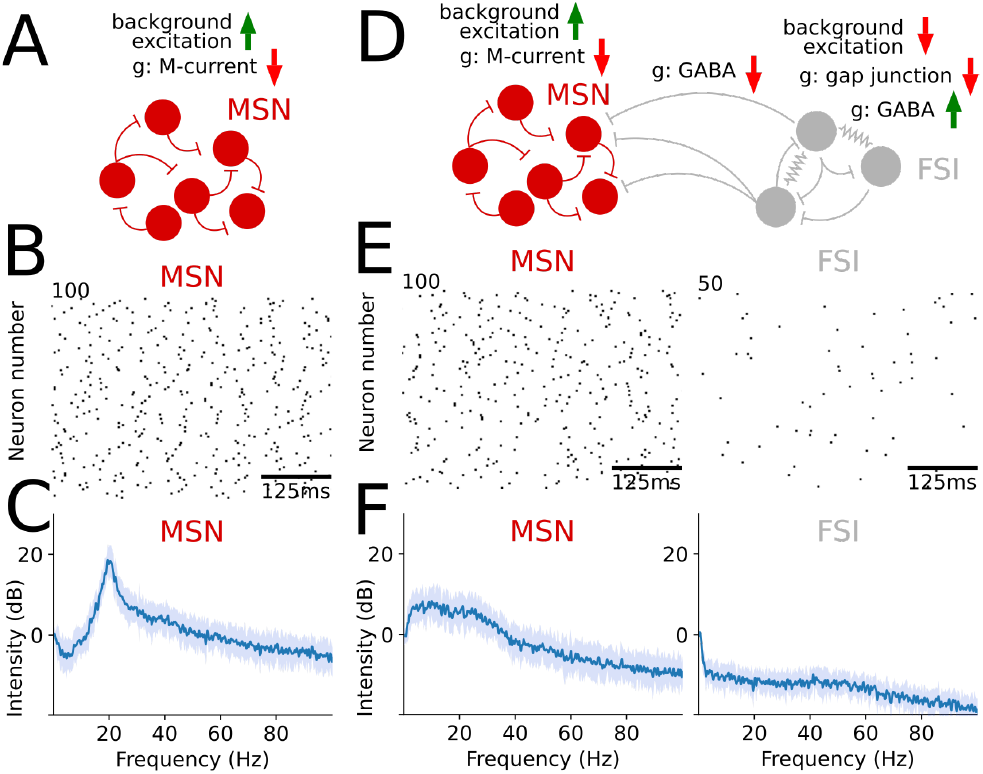
Population dynamics in PD, in the core striatal model. **(A)** Schematic illustrating the parametric changes in the MSN network during PD from parameters in baseline condition. **(B)** Raster plot showing spiking activity of MSNs during PD for the network in (A) (MSN firing rate: mean SD=7.66 0.04 spk.s^−1^, N=25 simulations). **(C)** Graphs showing the average (blue) and standard deviation (light blue) of the spectrum the MSN population activity, during PD for the network in (A) (N=25 simulations). **(D)** Schematic illustrating the changes in the parameters of the biophysical model during PD, from baseline condition, in the core striatal model. **(E)** Raster plots showing spiking activity of MSN and FSI neurons, all in PD for the network in (D) (MSN firing rate: mean SD=6.62 0.21 spk.s^−1^ N=25 simulations). **(F)** Graphs showing the average (blue) and standard deviation (light blue) of the spectra of MSN and FSI population activity, all in PD for the network in (D) (N=25 simulations).

**Figure S6:**
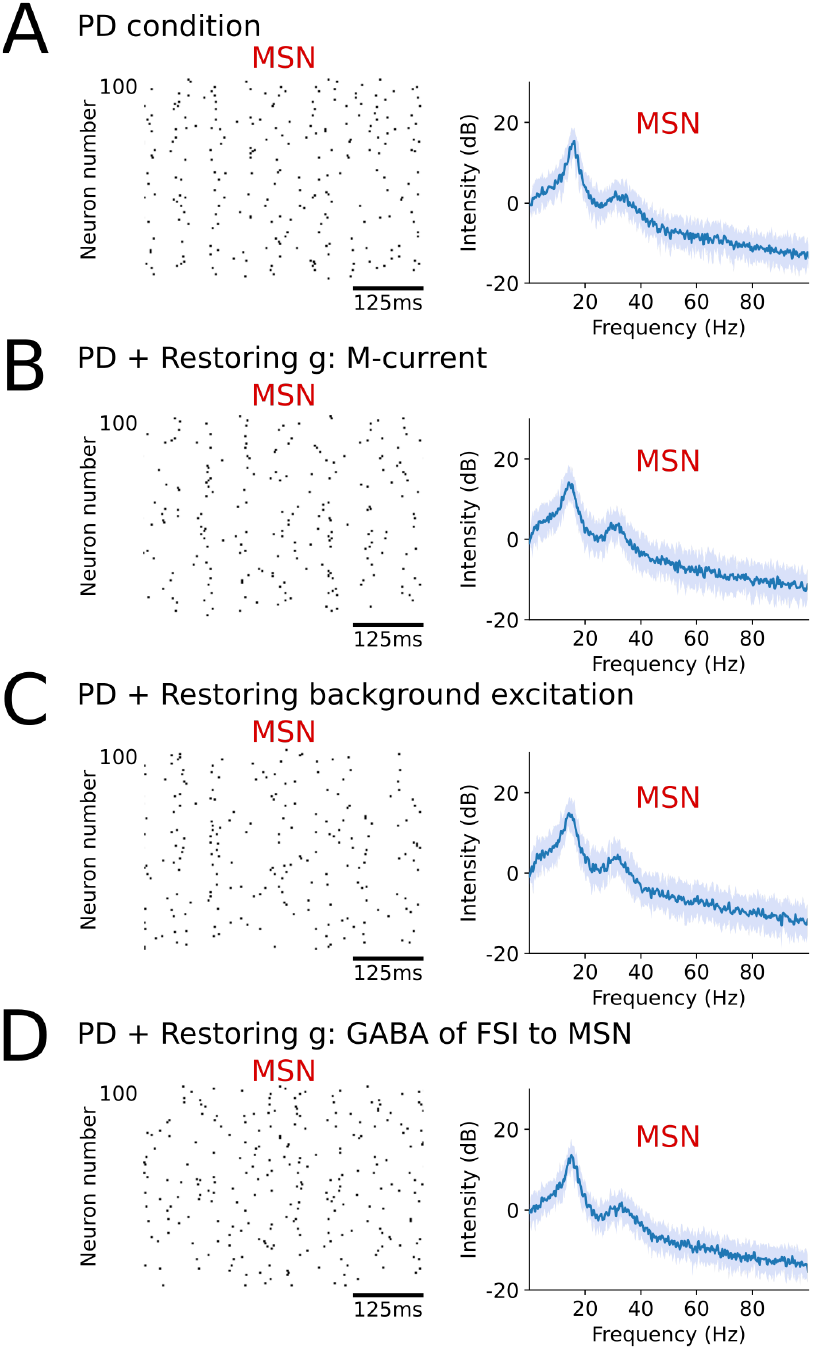
The effect of the biophysical perturbation in the MSN population on its activity during Parkinsonian conditions. **(A)** Left. Raster plot showing MSN spiking activity (MSN firing rate: mean±SD=4.85 ± 0.13 spk.s^−1^, N=25 simulations) during PD condition in the full network including FSIs, GPe and STN. Right. Graph showing the average (blue) and standard deviation (light blue) of the spectrum of MSN population activity during PD condition. **(B)** Similar to (A) but keeping the maximal conductance of the MSN M-currents intact during PD conditions (instead of decreasing it) (MSN firing rate: mean SD=3.99 0.10 spk.s^−1^, N=25 simulations). **(C)** Similar to (A) but keeping the background excitation onto MSNs intact during PD conditions (instead of increasing it) (MSN firing rate: mean SD=4.14 0.13 spk.s^−1^, N=25 simulations). **(D)** Similar to (A) but keeping the maximal conductance for the FSI projections to MSNs intact during PD conditions (instead of decreasing it) (MSN firing rate: mean±SD=4.73 ± 0.13 spk.s^−1^, N=25 simulations).

**Figure S7:**
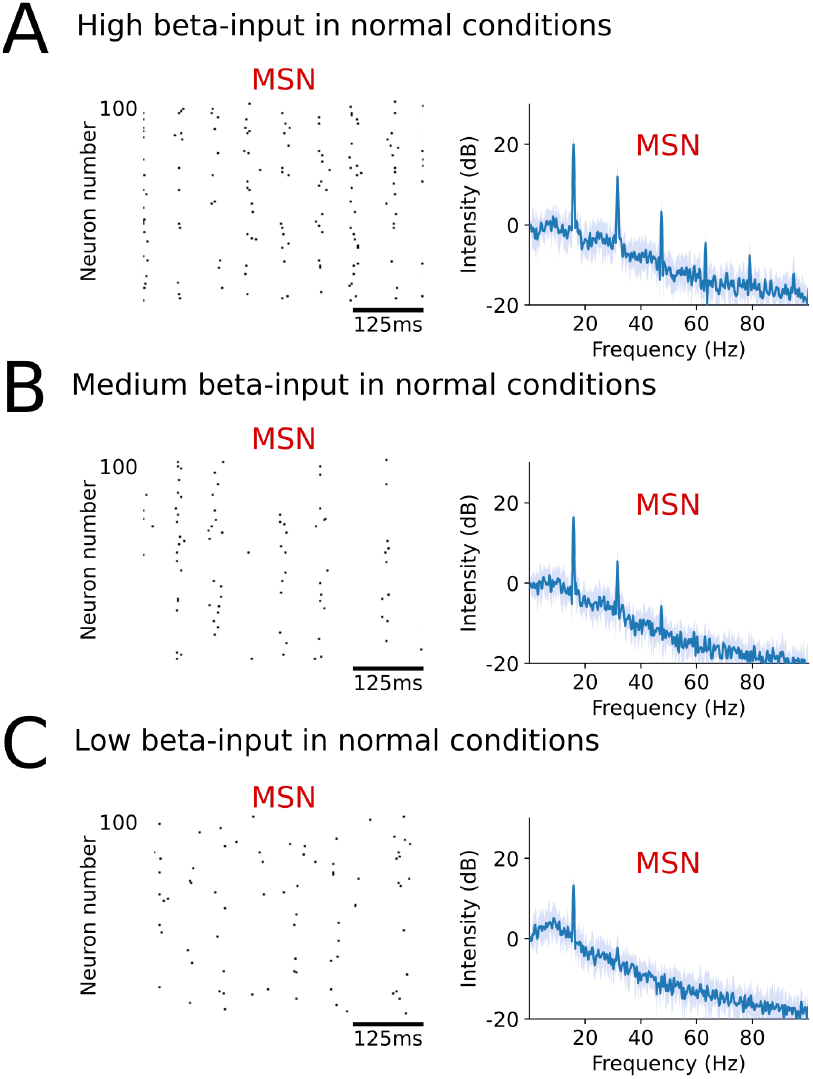
Beta activity in MSNs in baseline condition following exogenous beta-band input. **(A)** Left. Raster plot showing MSN spiking activity in baseline condition and a full network, with a high amplitude exogenous sinusoidal input current (at beta frequency of 15.8 Hz) applied to the MSNs (MSN firing rate: mean SD=3.24 0.18 spk.s^−1^, N=10 simulations). Right. Graphs showing the average (blue) and standard deviation (light blue) of the spectrum of MSN population activity, in baseline condition with a high amplitude exogenous sinusoidal input current applied to the MSNs. **(B)** Similar to (A) but decreasing the amplitude of the exogenous input into MSNs (MSN firing rate: mean SD=1.85 0.10 spk.s^−1^, N=10 simulations). **(C)** Similar to (B) but further decreasing the amplitude of the exogenous input into MSNs (MSN firing rate: mean±SD=1.34 ± 0.07 spk.s^−1^, N=10 simulations).

**Figure S8:**
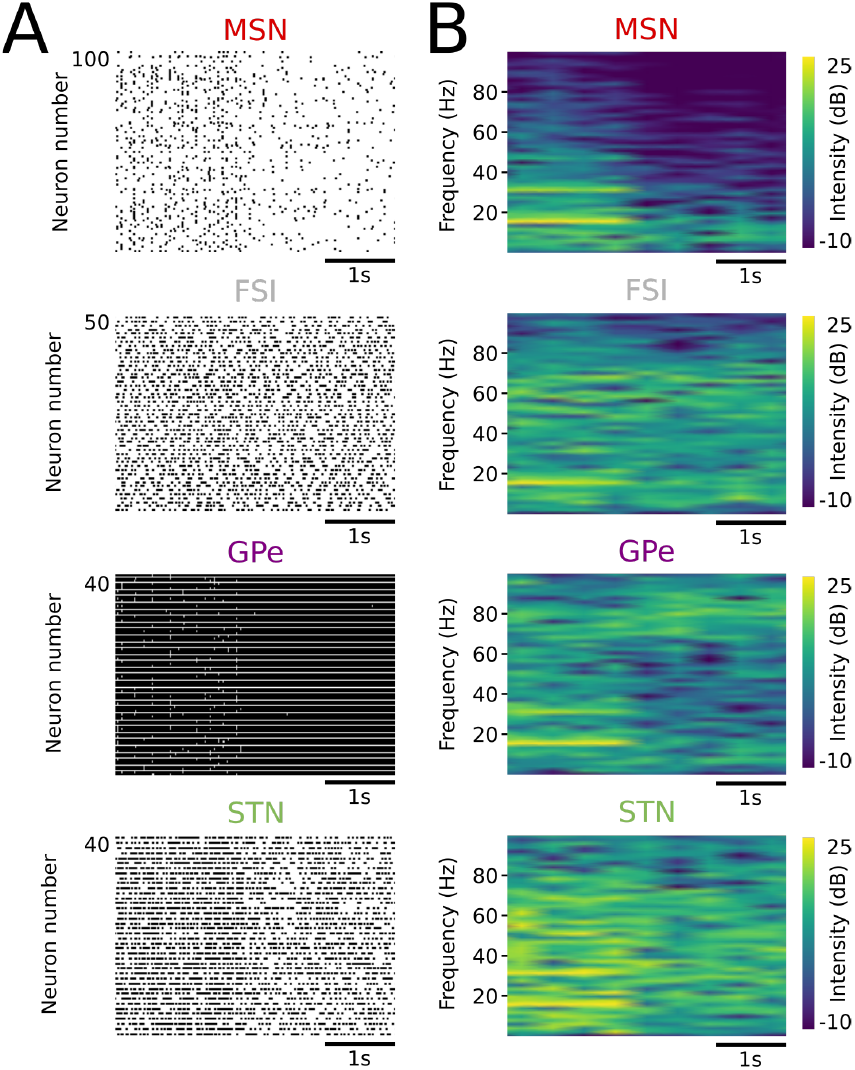
Resonance properties of the closed-loop system. **(A)** Raster plots showing spiking activity of the four neuronal populations in baseline condition, where a high amplitude exogenous sinusoidal (15.8 Hz) input current is applied to MSNs for the first 2 seconds of the plots, and then removed for the remaining 2 seconds. **(B)** Graphs showing the spectrogram of population activity for the four neuronal population, in the same conditions as in (A), where beta input is applied to MSNs for the first 2 seconds and then removed for the remaining 2 seconds.

**Figure S9:**
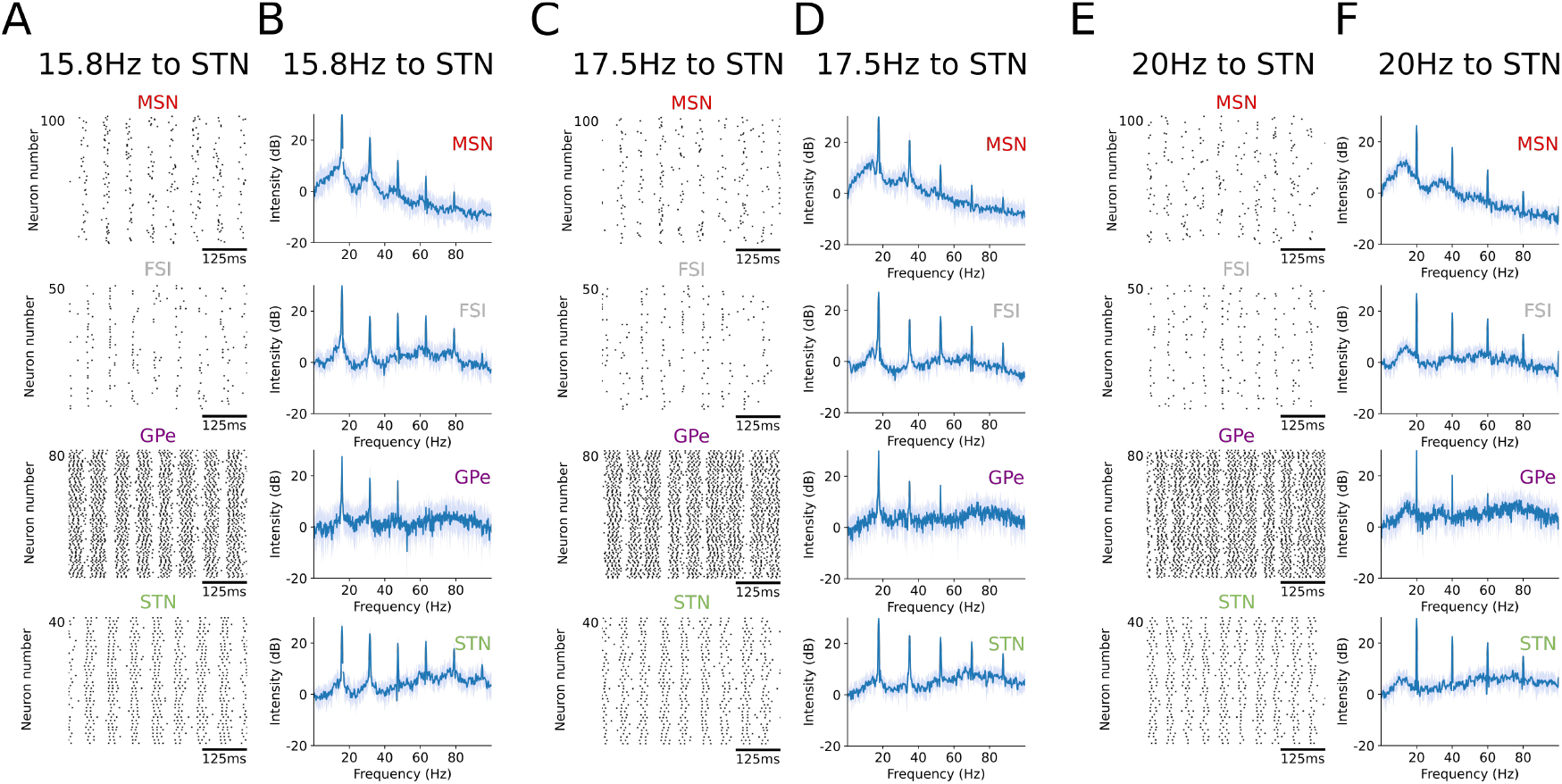
The effect of exogenous beta activity in STN during PD condition. **(A)** Raster plot showing spiking activity of the four populations in PD condition while an exogenous sinusoidal input current (at beta frequency of 15.8 Hz) is applied to STN (MSN firing rate: mean SD=5.83 0.12 spk.s^−1^, N=10 simulations). **(B)** Graphs showing the average (blue) and standard deviation (light blue) of the spectrum of population activity for each of the four populations, in PD condition while an exogenous sinusoidal input current is applied to STN (15.8 Hz) (N=10 simulations). **(C)** Same as (A) but with an input frequency of 17.5 Hz (MSN firing rate: mean SD=5.23 0.13 spk.s^−1^, N=10 simulations). **(D)** Same as (B) but with an input frequency of 17.5 Hz (N=10 simulations). **(E)** Same as (A) but with an input frequency of 20 Hz (MSN firing rate: mean SD=4.89 0.05 spk.s^−1^, N=10 simulations). **(F)** Same as (B) but with an input frequency of 20 Hz (N=10 simulations).

**Figure S10:**
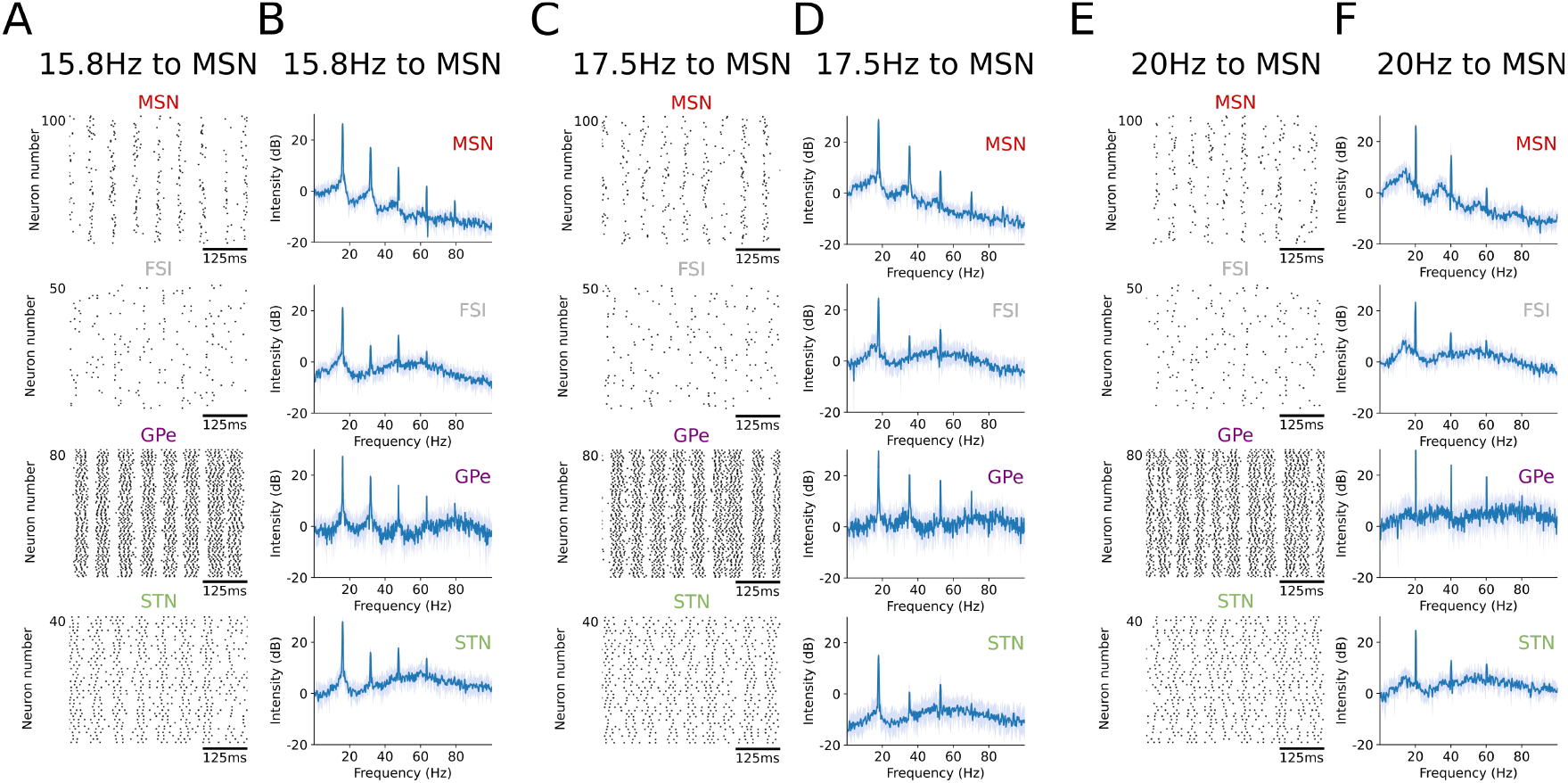
The effect of exogenous beta activity in MSNs during PD condition. **(A)** Raster plot showing spiking activity of the four populations in PD condition while an exogenous sinusoidal input current (at beta frequency of 15.8 Hz) is applied to MSN (MSN firing rate: mean SD=6.92 0.19 spk.s^−1^, N=10 simulations). **(B)** Graphs showing the average (blue) and standard deviation (light blue) of the spectrum of population activity for each of the four populations, in PD condition while an exogenous sinusoidal input current is applied to MSN (15.8 Hz) (N=10 simulations). **(C)** Same as (A) but with an input frequency of 17.5 Hz (MSN firing rate: mean SD=6.69 0.22 spk.s^−1^, N=10 simulations). **(D)** Same as (B) but with an input frequency of 17.5 Hz (N=10 simulations). **(E)** Same as (A) but with an input frequency of 20 Hz (MSN firing rate: mean SD=6.06 0.19 spk.s^−1^, N=10 simulations). **(F)** Same as (B) but with an input frequency of 20 Hz (N=10 simulations).

**Figure S11:**
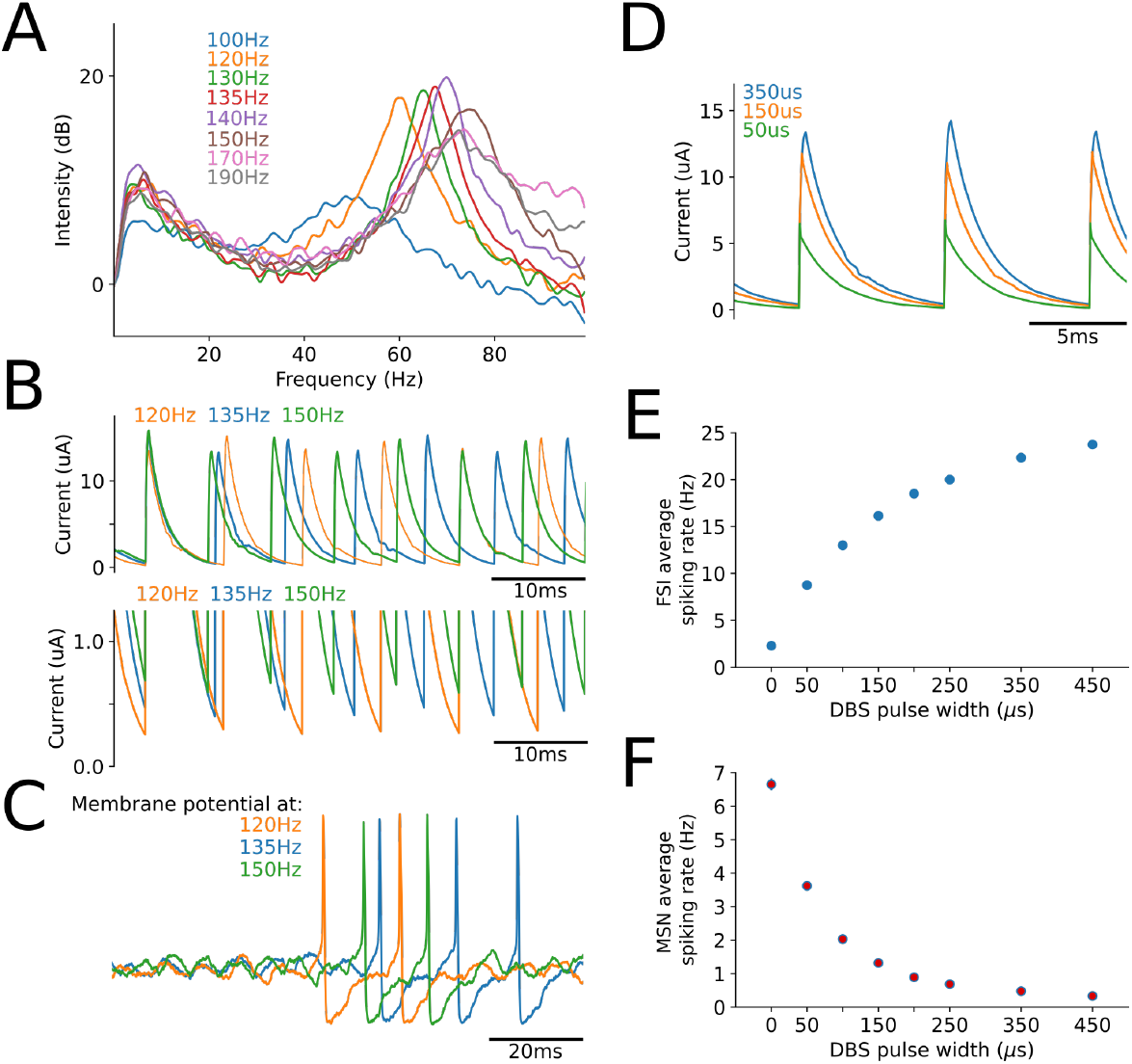
Effect of DBS frequency and pulse width on FSI and MSN activity during PD. **(A)** Plot showing the spectrum of FSI population activity for different stimulation frequencies. **(B)** Top. Plot showing the STN AMPA current, input into FSIs, for different stimulation frequencies. Bottom. Close-up, of the top plot, on the minima of the AMPA current. **(C)** Plot showing example traces of FSI membrane potentials for different stimulation frequencies. **(D)** Plot showing the STN AMPA current, input into FSIs, for different pulse widths. **(E)** Graph showing the average FSI spiking rate (vertical error bars represent standard deviation. The coefficient of variation std/mean is small for the error bars to be visible.) as a function of DBS pulse width (N=10 simulations for each value of pulse width). **(F)** Graph showing the average MSN spiking rate (vertical error bars represent standard deviation. The coefficient of variation std/mean is small for the error bars to be visible.) as a function of DBS pulse width (N=10 simulations for each value of pulse width).

**Figure S12:**
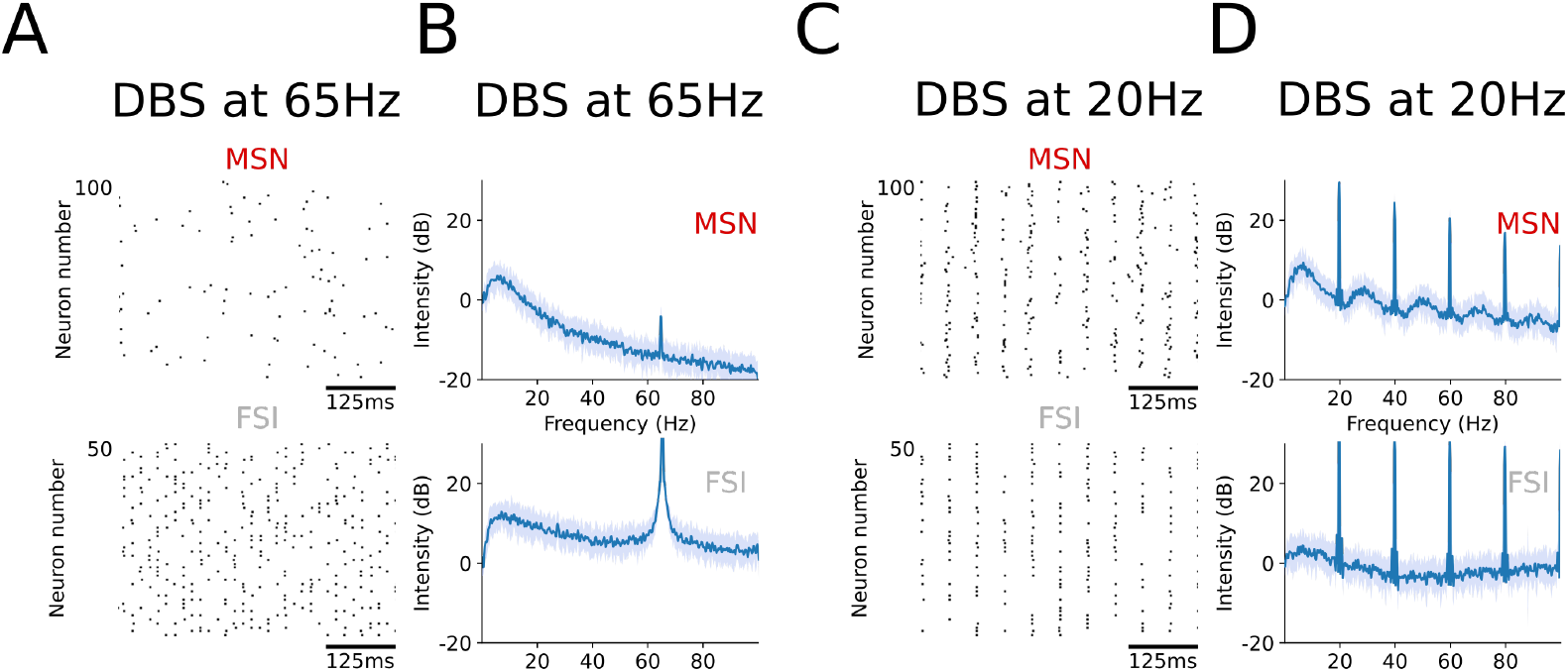
The effect of DBS with low frequencies. **(A)** Raster plot showing spiking activity of MSNs and FSI in PD condition while DBS is applied with a stimulation frequency of 20Hz (MSN firing rate: mean SD=5.57 0.23 spk.s^−1^, N=25 simulations). **(B)** Graphs showing the average (blue) and standard deviation (light blue) of the spectrum of population activity for MSNs and FSIs while DBS is applied with a stimulation frequency of 20Hz (N=25 simulations). **(C)** Same as (A) but with simulation frequency of 65Hz (MSN firing rate: mean SD=1.65 0.08 spk.s^−1^, N=25 simulations). **(D)** Same as (B) but with a stimulation frequency of 65Hz (N=25 simulations).

**Figure S13:**
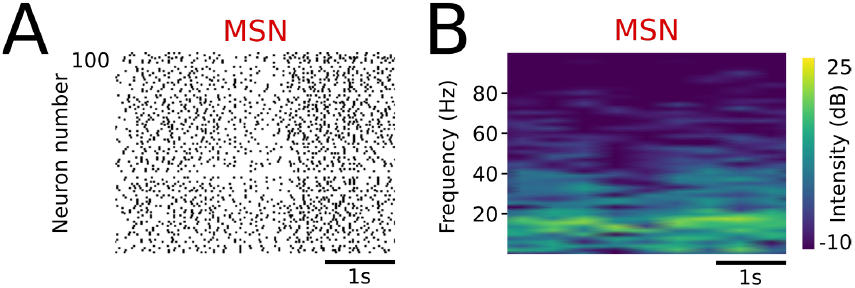
Variations in MSN beta activity through changes in MSN excitability during PD condition. **(A)** Raster plots showing spiking activity of MSNs during PD while excitability is varied through the applied current (*I_app_*), as a proxy for cortical/thalamic input. **(B)** Graphs showing the spectrogram of MSN population activity, in the same conditions as in (A).

**Figure S14:**
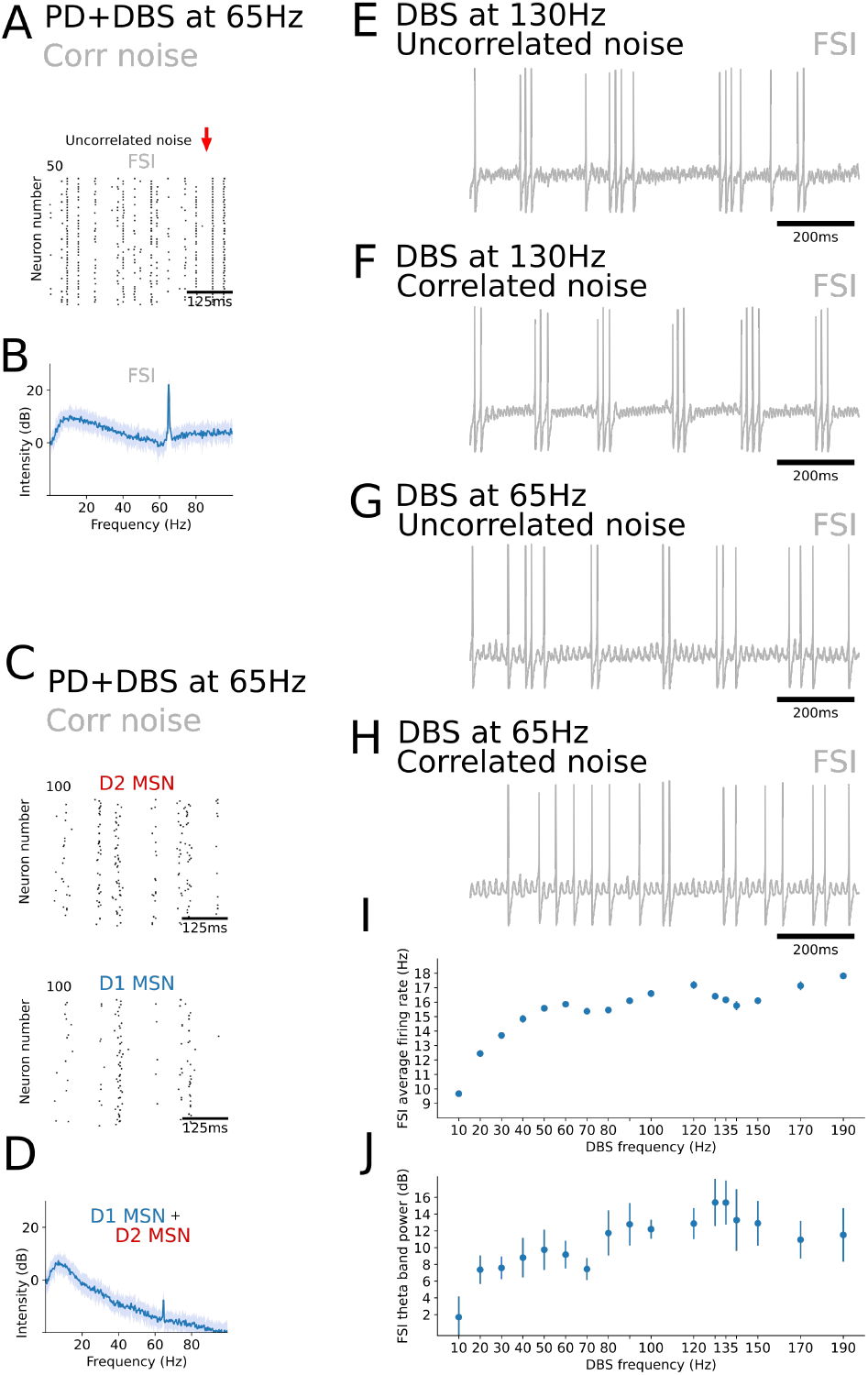
The effect of DBS with low frequencies under correlated noise. **(A)** Raster plot showing spiking activity of FSI neurons, in PD during DBS at 65Hz under correlated noise condition. **(B)** Graph showing the average (blue) and standard deviation (light blue) of the spectrum of FSI population activity, in PD during DBS at 65Hz under correlated noise condition. **(C)**, **(D)** Graphs displaying population activity of MSN neurons, as in (A) and (B), in PD during DBS at 65Hz under correlated noise. The raster plots show D1 and D2 MSN activity separately. **(E)** Graph showing the membrane potential of an example FSI neuron in PD during DBS at 130Hz under uncorrelated noise condition. **(F)** Same as (E) but during DBS at 130Hz under correlated noise condition. **(G)** Same as (E) but during DBS at 65Hz under uncorrelated noise condition. **(H)** Same as (E) but during DBS at 65Hz under correlated noise condition. **(I)** Graph showing the average FSI firing rate (bars representing standard deviation) as a function of DBS frequency (10 simulations per simulated frequency), under correlated noise condition. **(J)** Graph similar to (I) showing the average FSI theta band power as a function of DBS frequency. The theta band power is an average of the intensity (dB) of the spectrum in the frequency interval 4-8Hz. The baseline power considered to compute the intensity is the average power in the band less than 1Hz.

